# Synaptic Development of Fine Spatial Scale Organization of Neuronal Orientation Tuning in Mouse Primary Visual Cortex

**DOI:** 10.64898/2026.07.16.738755

**Authors:** Peijia Yu, Gengshuo John Tian, Brent Doiron

## Abstract

Primary sensory cortices often organize neurons with similar stimulus preference into spatially functional maps. Recent work in mouse primary visual cortex (V1) has established that neuronal tuning to the orientation of visual grating stimuli is organized into ‘micro-clusters’, where physically close neuron pairs (~ 20 *µ*m) share highly similar orientation preferences, but the organization is unstructured beyond this narrow range. This fine-scale organization is seemingly at odds with the underlying intracortical circuitry in mouse V1 whose spatial extent is an order of magnitude broader (100 ~ 200 *µ*m). In this study, we explore an activity-dependent synaptic plasticity model of spatially structured thalamo-cortical connectivity. We develop theory under asymptotic conditions specific for mouse V1, and derive concrete circuit conditions under which ‘micro-clusters’ naturally develop. In particular, the recurrent interaction among V1 neurons requires an additional component over a ‘micro’-spatial scale, while the spatial profiles of balanced excitation and inhibition support an effective ‘micro’-scale interaction. Together, our results provide a developmental mechanism and analytical framework linking thalamo-cortical development, recurrent circuit structure, and the emergence of functional organization in primary visual cortex.

## I. INTRODUCTION

The rules that govern the organization of a biological system at one spatial scale, may not be in effect at other scales [1]. From protein [2] to ecological [3, 4] networks, how interactions between components differ across spatial scales contribute to the rich and complex spatial patterns in system organization. The nervous system is a particularly compelling example, as operative spatial scales span six orders of magnitude [5], with distinct interaction rules for subcellular networks [6] all the way to whole brain circuits [7, 8].

A universal organizing principle of synaptic connectivity in the neocortex is the ‘spatial dependent rule’: pairs of neurons that are closer to each other have a higher likelihood of having direct synaptic connections [9–13]. Such local wiring is thought to support the pinwheel-like columnar structure of orientation tuning preference, where neighboring neurons tend to share similar tuning preferences, that is reported in the visual systems of the cat [14], ferret [15], tree shrew [16] and several non-human primate species [17, 18]. By contrast, the primary visual cortex (V1) of rodents appears to lack any clear spatial organization of orientation tuning preference, often termed as ‘salt-and-pepper’ structure [19– This disordered representation is in apparent opposition to the well-established spatial dependence of the intracortical synaptic connectivity in mouse V1 [12, 13].

Despite these original reports of disordered structure in mouse V1, recent population imaging studies have provided conflicting evidence of measurable spatial clustering of tuning in mouse V1 over fine spatial scales [23– In our prior study [26], using a refined experimental approach we identified that neurons in both layer 2/3 (L2/3) and layer 4 (L4) of mouse V1 form strong spatial clusters of similarly tuned neurons over an extremely *narrow* spatial scale (~ 20 *µ*m), while being randomly organized on *broad* spatial scales (> 30 *µ*m) – an organization we termed ‘microclustered’ (Fig. 1). Micro-clusters are far narrower than the commonly reported spatial scale of intracortical recurrent connection footprints in mouse V1, which typically span on the order of 100 ~ 200 *µ*m [9, 11–13] This mismatch raises a central question: how could spatially broad circuitry give rise to a functional organization that is expressed on a narrow spatial scale?

**FIG. 1.**
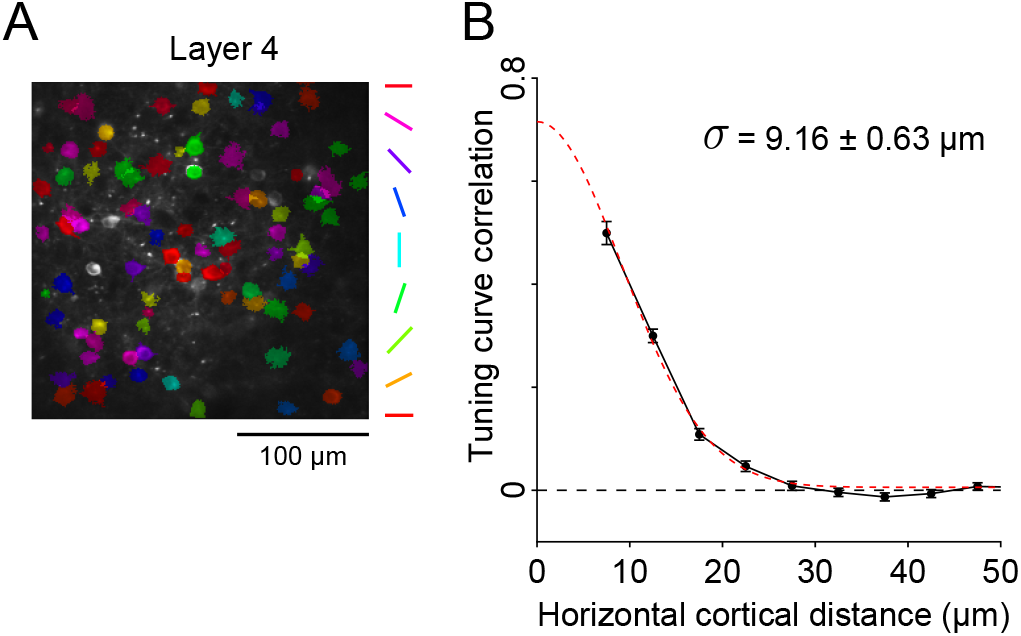
‘Micro-cluster’ organization of orientation tuning in L4 of mouse V1. (A) Example *in vivo* fluorescence image of GCaMP6^+^ L4 neurons. Colors represent the preferred orientation of neurons. (B) A distinguishing feature of ‘micro-cluster’ organization: the correlation between tuning responses of pairs of neurons sharply drops as a function of the horizontal cortical distance between neurons. Error bars represent standard error of the mean. The red dashed curve is the Gaussian fitting, where *σ* represents the standard deviation / spatial width (error represents the 95% confidence interval). The Figure is a reproduction of the central results in [26].

In the canonical pathway of early vision, information starts from the retina and is conveyed by the lateral geniculate nucleus (LGN) to neurons in layer 4 (L4) of V1 [14]. Neurons in L2/3 are mostly driven by L4 activity, rather than directly from LGN (i.e., LGN → L4 → L2/3) [27]. Thus, the measurement of micro-clusters in both L4 and L2/3 of mouse V1 [26] creates two interlinked questions: (1) How does micro-clustered functional organization emerge in L4 from thalamic input? (2) How is micro-cluster organization propagated to L2/3 from L4 through intracortical interactions? Our previous study [26] addressed the second problem by predicting that reconciling the fine spatial scale of the functional map with circuitry constraints requires two circuit features. First, a previously underappreciated, additional component of intracortical recurrent synaptic wiring that operates at a similar fine spatial scales to that of micro-cluster. Second, an effective excitation-inhibition (E-I) balance of intracortical recurrent connectivity over the known, ‘macro’-spatial scale, but an E-I imbalance over the fine, ‘micro’-spatial scale. These hypotheses were tested through various complementary experimental investigations of connectivity mapping of L2/3 → L2/3 and L4 → L2/3 in mouse V1 [26, 28].

In this study, we address the first problem: the emergence of a micro-clustered organization of orientation preference in L4 neurons of mouse V1. A natural hypothesis is that it arises during the development process through activity-dependent plasticity of LGN-L4 synapses before eye opening, which has been proposed for the codevelopment of orientation-selective receptive fields [29, 30] and pinwheel-like orientation map together in cat or primate V1 [31]. Yet, within the framework of LGN-L4 plasticity, the mechanics underlying the fine spatial scale of micro-clusters remains unclear (but see [32] for a non-plasticity-based model).

Previous studies have formulated models of Hebbian development as a self-organization process [33–36]. We employ a similar framework for the plasticity of thalamo-cortical synapses from LGN to L4 within a spatially structured cortical network, building on a seminal past study [31]. We develop analytic theory for the principal eigen-values and eigenfunctions of the linear operator associated with Oja’s learning rule [37, 38], under asymptotic conditions specific for the physiological constraints of mouse V1. In this framework, orientation-selective receptive fields admit a compact description in terms of circular-harmonic basis functions. We find that the emergence of a micro-cluster organization in L4 requires qualitatively similar circuit constraints to those underlying the propagation problem in our previous study [26], namely an additional micro-spatial scale component of the L4 → L4 recurrent interaction together with effective macro-scale E-I balance that suppresses long-range pattern formation and a micro-scale E-I imbalance that permits fine-scale structure. Additionally, the spatial scale of correlations in the activity of LGN units also impacts the spatial scale of micro-cluster development.

In total our work, combined with our past study [26], provides a novel theory of the emergence of a narrow spatial scale of tuning organization in mouse V1, while ensuring a lack of organization over broad spatial scales.

## II. RESULTS

### A. Framework of the network model in Oja’s learning rule

In this section we present our learning model that is adapted from the classic model presented in [31]. To study the emergence of functional organization of orientation tuning preference in layer 4 (L4) of mouse V1 through thalamo-cortical synaptic learning, we consider a network of LGN units and L4 neurons on respective two-dimensional square lattices (Fig. 2A; also see Methods for details). The field 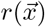 denotes the activity of a L4 neuron at location 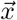, while 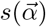 gives the activity of a LGN unit at location 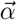. In this network L4 neurons receive inputs from LGN units through feedforward connections, and from other L4 neurons through intracortical recurrent connections. As a simplification, the subtypes of ON or OFF LGN units, and excitatory or inhibitory neurons, are not explicitly modeled. The strength of feedforward connection to L4 neuron at 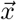 from LGN unit at 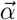 is given by 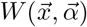, and the intracortical recurrent connection to L4 neuron at 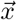 from another neuron at 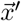 is given by 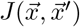 (Fig. 2A). We assume the strength of intracortical recurrent connections only depend on the separation distance between neurons, i.e. 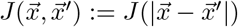.

**FIG. 2.**
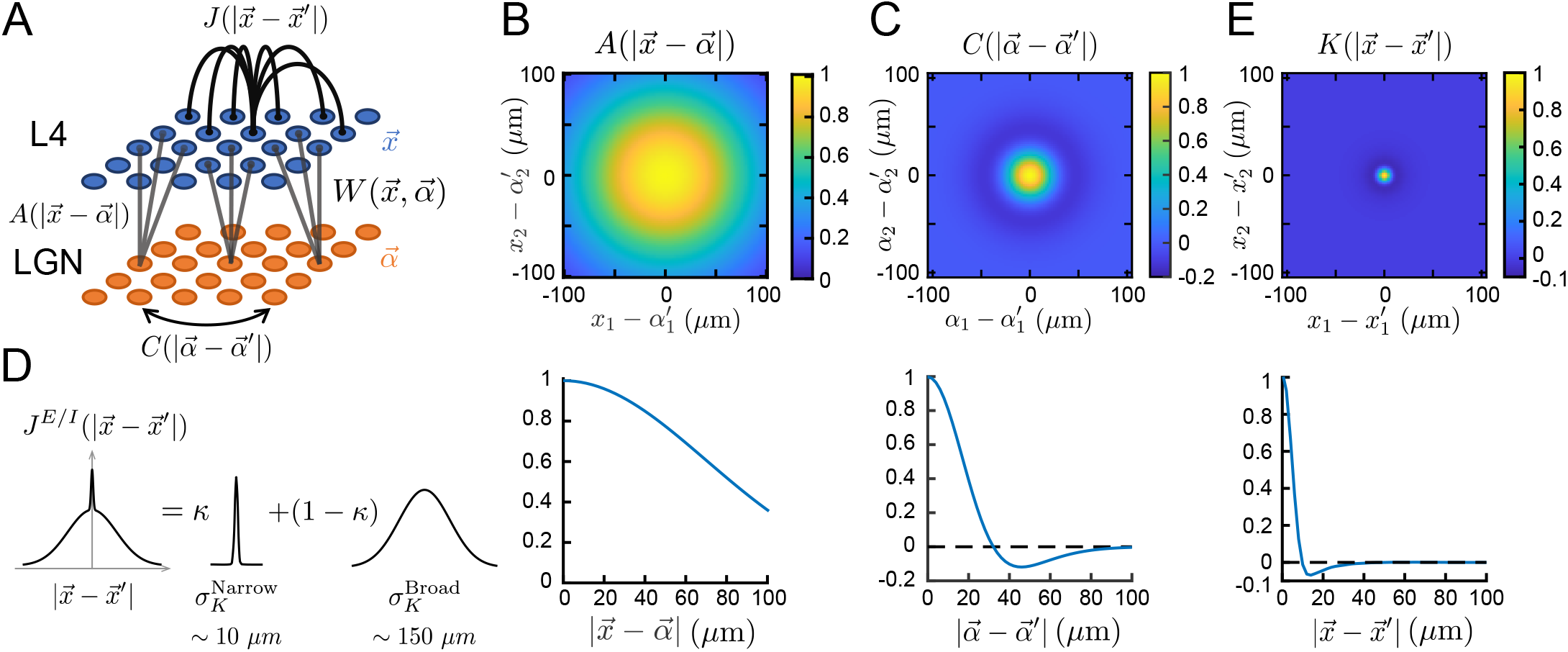
The model framework and main components of the learning dynamics of network model. (A) Schematic of two-layer network model for LGN and L4 of mouse V1. 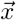 and 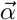 denote the respective positions of L4 neurons and LGN units on two-dimensional grids. 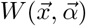 represent the thalamo-cortical connection strength, which is plastic for learning process. 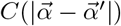 represents the correlation function of activity of LGN units. 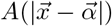 represents the arbor function that restricts the thalamo-cortical connection to be spatially convergent. Finally, 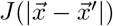 represents the intracortical recurrent connection strength among L4 neurons;. The functions *C, A* and *J* are assumed to be non-plastic for simplicity. (B) The spatial profile of the arbor function 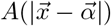. The top panel gives the full 2-dimensional profile, while the bottom panel gives the radial dependency. (C) The spatial profile of the correlation function between LGN unit activity 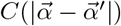. (D) The intracortical recurrent connection strength 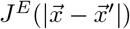 or 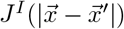 are modeled as sum-of-Gaussians function, with one component over *narrow* spatial scale (characterized by 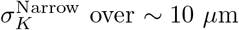) with ratio *κ*, and another component over *broad* spatial scale (characterized by 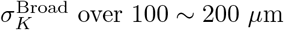) with ratio 1 − *κ*. (E) The spatial profile of effective intracortical recurrent interaction function 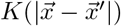.

#### A.1 Network Dynamics

The dynamics of L4 neuron responses obey the following linear system:

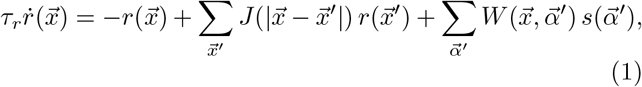

where *τ*_*r*_ is an effective time constant. Assuming periodic boundary conditions on the cortical lattice, the steady state solution has the form (see Appendix A for details):

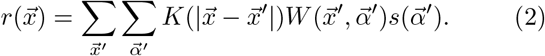

Here 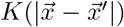 is the effective convolution kernel representing intracortical recurrent connections, which is determined by 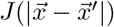 in Fourier space (see Appendix A for details):

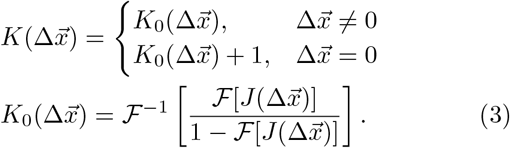

#### A.2 Synaptic Plasticity

To simplify our analysis we take the effective intracortical recurrent interaction 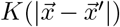 to be non-plastic, and only model the plasticity of the feedforward synaptic connections, 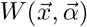. These connections obey the following dynamics:

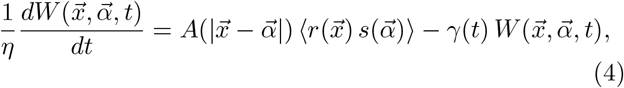

where *η* is the learning rate (see Methods for details). The function 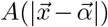 models the arborization of axonal LGN inputs from different locations 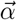 in the receptive field to L4 neuron at location 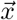. We assume the arbor function to be non-plastic, and only depends on their separation distance 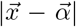 (Fig. 2A). The expression 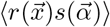 represents how the Hebbian learning rule is driven by the covariance between L4 neurons and LGN unit activity, with ⟨·⟩ being the temporal average. Finally, *γ*(*t*) is a time-varying normalization factor that scales 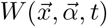 to regularize the Hebbian learning dynamics. This form is such that the Frobenius norm of 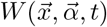 is constant over time [37, 39]:

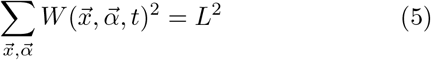

(see Methods for details of L).

Since the dynamics of L4 neuron responses attain steady state in a much faster timescale than the learning process, we substitute the steady state solution of neuron responses (Eq. (2)) into the synaptic learning equation (Eq. (4)), to yield the dynamics of 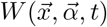 following Oja’s learning rule [37, 38]:

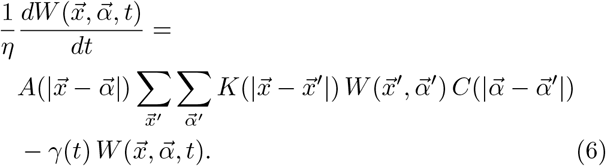

Here the term

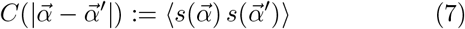

represents the correlation function between the activities of LGN units that are separated by distance 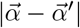(Fig. 2A).

Eq. (6) is the core learning model of our study, and we seek to determine the conditions for its solution to promote a micro-clustered organization of orientation preference in L4. To do so we first need to specify the spatial scales of the various circuit components that impact learning dynamics.

#### A.3 Network Spatial Scales

The learning dynamics given in Eq. (6) contain three spatial kernels: 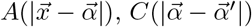 and 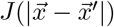.

The arbor function 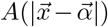 is taken to be a Gaussian function (Fig. 2B):

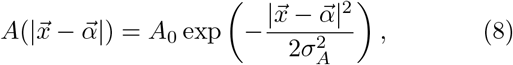

where *A*_0_ and *σ*_*A*_ represents the amplitude and spatial spread, respectively (see Methods for details). Note that *σ*_*A*_ in mouse V1 is large so that the arbor function covers a broad range in physical space [21].

The LGN correlation function 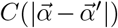 describes the correlation of spontaneous LGN activity before eye opening that facilitates the development of orientation selectivity and its spatial organization in mouse V1. This can be well described by a ‘Mexican-hat’ difference-of-Gaussians function when both ON-center and OFF-center receptive fields are taken into account [31] (Fig. 2C; see Methods for details):

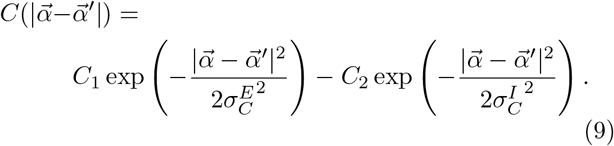

The intracortical recurrent connection strength between L4 neurons, 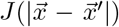, is guided by past experimental studies on mouse V1 cortical circuits that report spatial scales spanning 100 ~ 200 *µ*m [9–13]. The clear broad spatial scales of 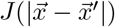 lead to a mechanistic puzzle: how is it that mouse V1 neurons form a micro-clustered functional organization of tuning preference over a *narrow* spatial scale over ~ 10 *µ*m (Fig. 1) while the underlying intracortical recurrent circuitry spreads over *broad* spatial scales over 100 ~ 200 *µ*m? To resolve this apparent mismatch between *broad* circuitry and *narrow* functional organization, we follow our past work [26, 28] and assume that the spatial profile of intracortical recurrent connections contains an additional, previously underappreciated component that operates at *narrow* spatial scales comparable to micro-clusters over ~ 10 *µ*m. Specifically, we postulate that the spatial profile of both excitatory and inhibitory intracortical recurrent connection strength, 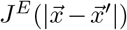 and 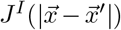, has following form of a sum-of-Gaussians functions (Fig. 2E):

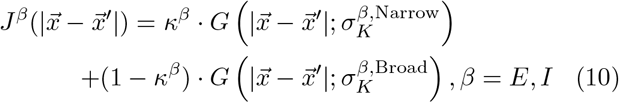

where

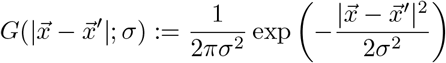

is the normalized two-dimensional Gaussian function. Here 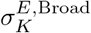 and 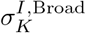 represent the spatial scales of *broad* Gaussian components over 100 ~ 200 *µ*m, while 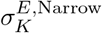 and 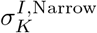 represent the spatial scales of *narrow* Gaussian components over ~ 10 *µ*m that are comparable to the spatial scales of micro-cluster organization. The coefficients *κ*^*E*^ and *κ*^*I*^ are the ratios of the narrow component to the overall connection strength, and 1 − *κ*^*E*^ and 1 − *κ*^*I*^ represent the ratios of the remaining broad components. In our model *κ*^*E*^, *κ*^*I*^ ≪ 1; this is consistent with our previous estimates [26, 28]. Finally, as a simplification we consider the overall intracortical recurrent connection among L4 neurons 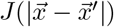 to be

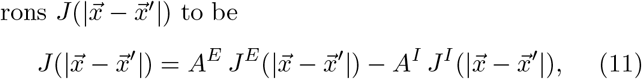

where *A*^*E*^ and *A*^*I*^ represent the relative strength of excitatory and inhibitory connections. Then 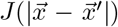 determines the effective intracortical recurrent interaction 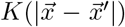 as Eq. (3). With appropriate choices of parameters *A*^*E*^ and *A*^*I*^, the excitatory and inhibitory interactions are mostly balanced (canceled) over broad spatial scales, while imbalanced over narrow spatial scales due to non-zero *κ*^*E*^ and *κ*^*I*^ (see Methods for details). This combination leads to an effective total interaction dominated by the narrow spatial scale only (Fig. 2E and [26]) that is much narrower than the LGN correlation function (Fig. 2C vs. Fig. 2E).

### B. Simulation: Emergence of orientation selectivity and micro-clustered organization

Numerical simulations of the learning model show Gabor function-like receptive fields gradually emerging over time (Fig. 3A, comparing *t* = 1000 and *t* = 8000 time points). To quantify the orientation selectivity of these receptive fields, we consider a grating visual stimulus in the LGN layer with the following form:

**FIG. 3.**
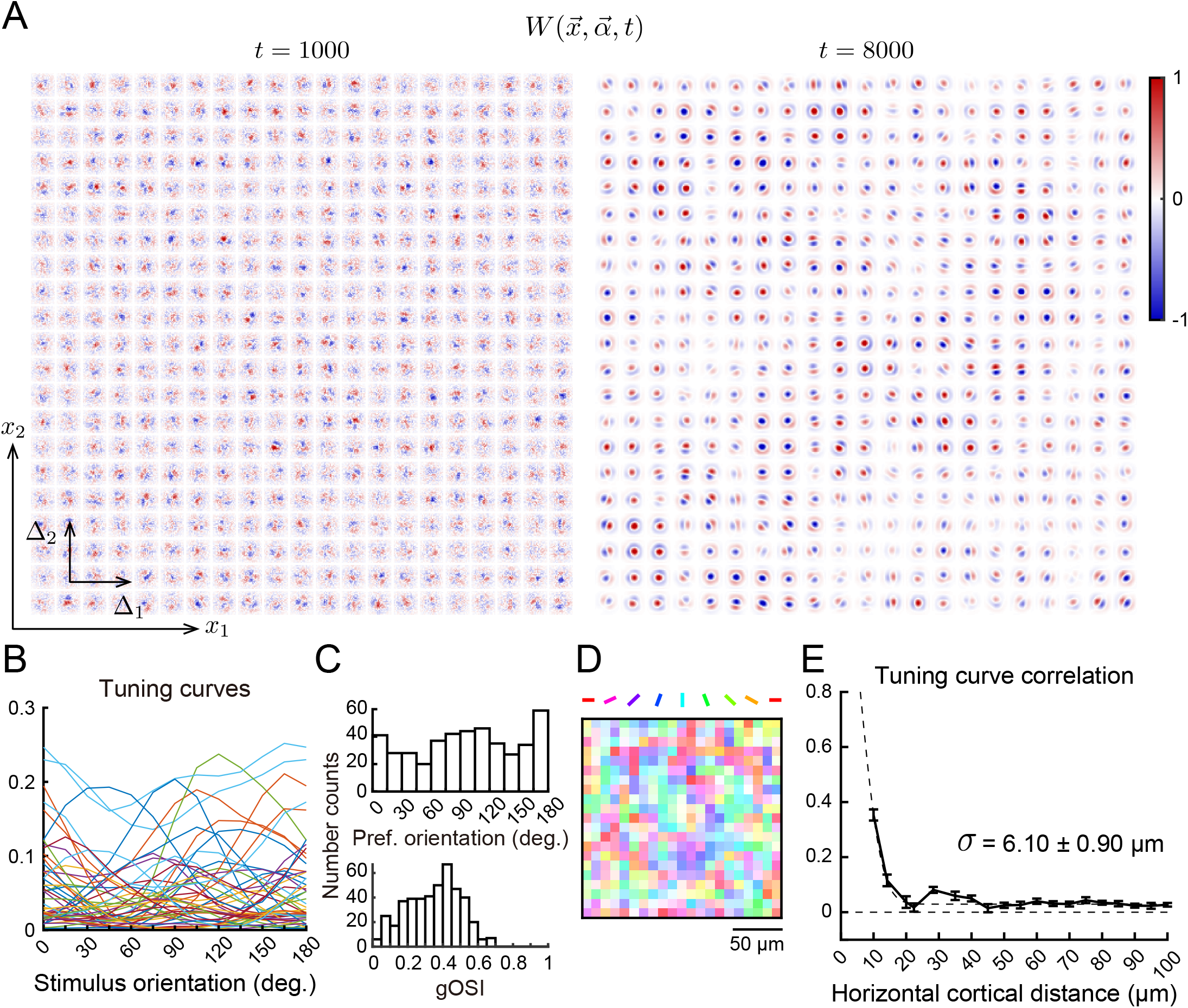
Emergence of receptive fields, orientation tuning selectivity and micro-cluster organization of orientation tuning preference. (A) The thalamo-cortical connection weight 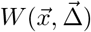 (where 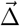 is translated receptive field location 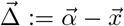) at an early time point (*t* = 1000) and a later time point (*t* = 8000) of learning process. Each small squared panel correspond to the specific position 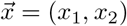 of different cortical neurons; within each panel is *W* over location 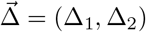. (B) Tuning curve responses of 50 random example neurons, at *t* = 8000 (also for all following panels). (C) Histogram distribution of the preferred orientations (above) and orientation selectivity index (gOSI) of L4 neuron population (below). (D) The preferred orientation map of L4 neuron population. (E) The correlation of tuning responses as a function of the distance between L4 neuron pairs. Error bars: standard error of mean; Dashed curve: fitting to Gaussian function; *σ*: the standard deviation / spatial width of fitting (error represents the 95% confidence interval).

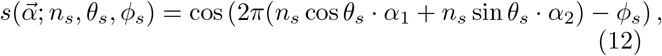

where *n, θ*_*s*_, and *ϕ*_*s*_ represent the spatial frequency, orientation and phase of the grating stimulus, respectively. The response of L4 neurons to this stimulus is defined as:

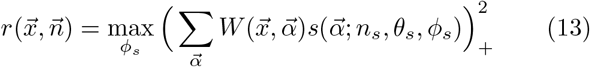

(where 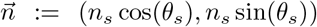), i.e. the rectified quadratic of the inner product of the receptive field and grating stimulus, maximized over stimulus phase (see Appendix F for details). In the following analysis we take the average over all responsive spatial frequencies *n* for the orientation tuning responses.

With standard parameters (see Methods for details) and at the final time point of the development process, robust orientation selectivity emerges in neuron population activity (Fig. 3B, 3C). Importantly, the correlation of orientation tuning responses as a function of distance between pair of neurons shows significantly strong correlation over close distances (~ 10 *µ*m), but then sharply asymptotes to near zero at farther distances (Fig. 3E). This corresponds to the micro-cluster organization observed in the experimental data (Fig. 1; also for the orientation preference map in Fig. 3D). These numerical results suggest that our model framework has the sufficient components to allow the emergence of micro-cluster organization of orientation selectivity through a biologically plausible learning process.

Our central goal is to understand how the spatial scale of tuning correlation as a function of distance (*σ* in Figure 3E) is shaped by the spatial scales of the arbor function 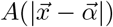, the LGN correlation function 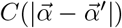, and the effective intracortical recurrent interaction 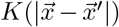, respectively. To this end, we first varied the learning model over different scaling factors applied to the spatial widths of these three functions, as *l*_*A*_, *l*_*C*_, and *l*_*K*_ (Fig. 4A). The spatial scale of tuning correlation over distance is significantly affected by the width of effective intracortical recurrent interaction 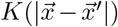 (Fig. 4B, 4C, along columns) and the LGN correlation function 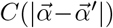 (Fig. 4B, along rows), while the arbor function 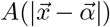 has negligible effect on the spatial organization of L4 tuning (Fig. 4C, 4D, along rows). It is important to note that maintaining a narrow spatial scale of 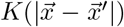 is necessary for the formation of micro-cluster over ~10 *µ*m (Fig. 4B, 4C, compare with the red box along columns).

**FIG. 4.**
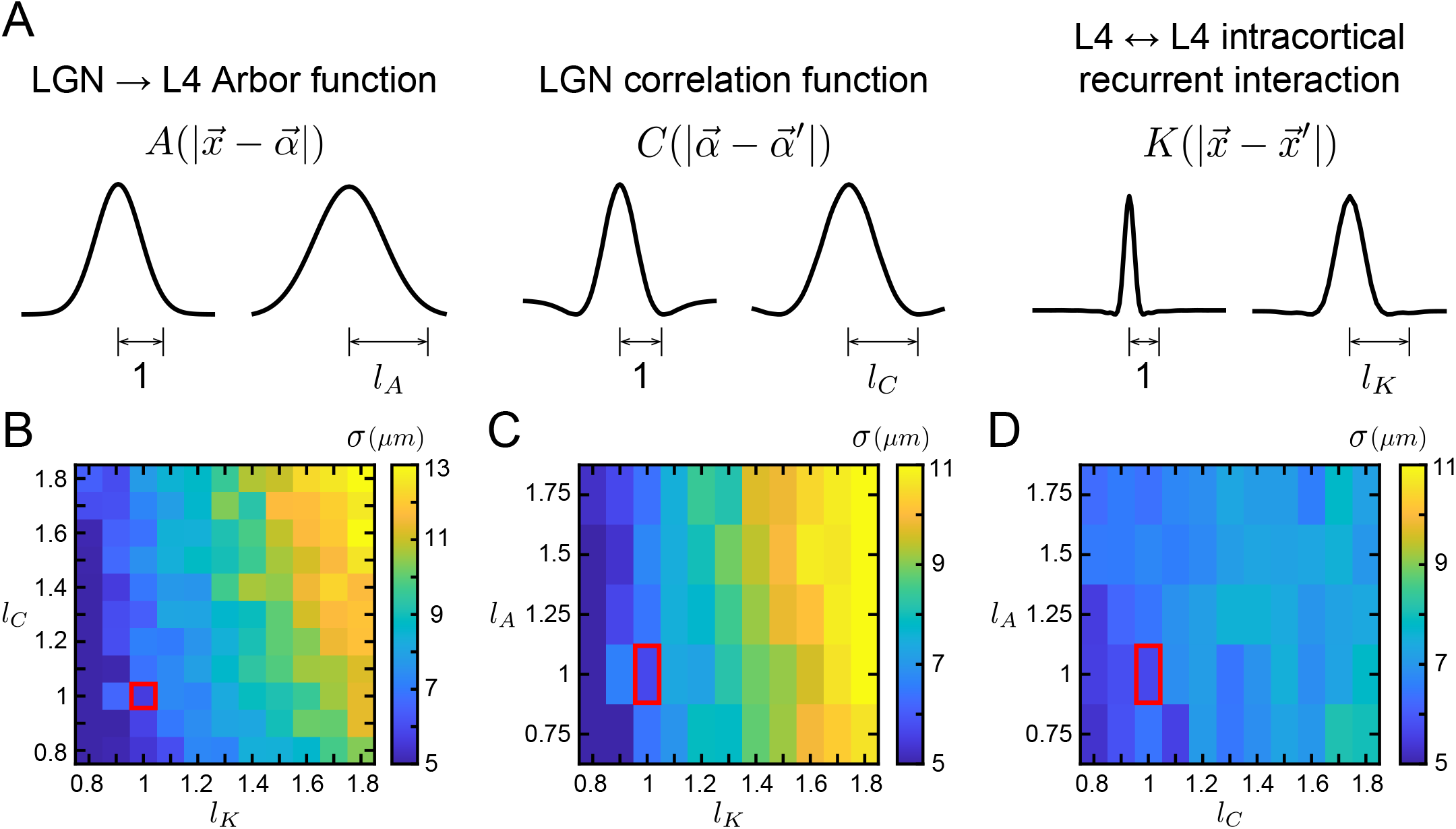
Determinants of the spatial scale of the ‘micro-cluster’ organization of orientation tuning preferences. (A) Schematic of *l*_*A*_, *l*_*K*_ and *l*_*C*_: the scaling factor of spatial scale of functions 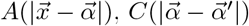 and 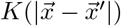. (B) The output spatial width *σ* of tuning correlation as a function of distance (same definition as in Figure 1B and Figure 3E), with different combinations of (*l*_*K*_, *l*_*C*_) in numerical simulations. Both *l*_*K*_ and *l*_*C*_ significantly affects *σ*. (*l*_*K*_, *l*_*C*_) = (1, 1) is marked by the red box as the standard parameter in Figure 2, 3. (C, D) Similar to (B) but with different combinations of (*l*_*K*_, *l*_*A*_) or (*l*_*C*_, *l*_*A*_). *l*_*A*_ has negligible effect on *σ*. (*l*_*K*_, *l*_*C*_) or (*l*_*C*_, *l*_*A*_) = (1, 1) are marked by red boxes as the standard parameter in Figs. 2, 3.

The strong dependencies of the learned micro-cluster organization on spatial scales *l*_*K*_ and *l*_*C*_, combined with its lack of dependence on *l*_*A*_, are important features of the learning process that we wish to understand. To this end, we next build a theory for the analytical solution of Eq. (6) under reasonable circuit assumptions.

### C. Asymptotic analytical solution of the leading eigenfunctions and eigenvalues of the learning equation

We first rewrite the continuous, integral version of the learning equation following Oja’s rule (Eq. (6)):

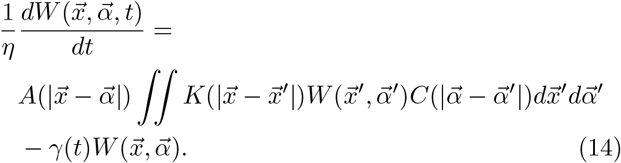

Previous studies have shown that Oja’s learning rule performs principal component analysis ([37–40]). Namely, the principal eigenfunctions that correspond to the largest eigenvalue *λ* of the linear operator of the Hebbian rule determine the final receptive fields after learning process:

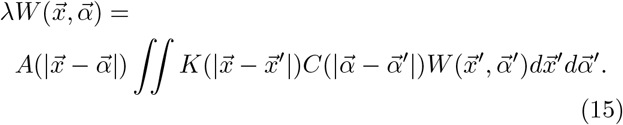

For this linear (Fredholm) integral equation, it is difficult to obtain a general analytic solution (excluding the trivial solution 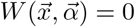 for all 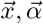), unless we restrict ourselves to the following asymptotic conditions (which are oversimplified for the physiological conditions of V1):

i. The limiting cases where any of the *A, C* or *K* functions are *δ*−functions or constants [41].
ii. *A, C* or *K* are all Gaussian functions [36].

To obtain a solution under proper asymptotic conditions for mouse V1 physiology, we will proceed in six sequential steps, organized into successive sections. Subsection C.1 will formally set the asymptotic conditions on the circuit and LGN activity as assumptions in the Fourier domain. Subsection C.2 will derive the eigensystem for 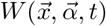 when the solution is assumed to be separated into radial and angular components. Subsection C.3 will solve the radial part of the eigenfunction. Subsection C.4 will solve the angular part of the eigenfunction, and determine the spatial frequency of principal eigenfunctions corresponding to the maximal eigenvalue. Subsection C.5 will compute the analytical form of the principal eigenfunctions back in the spatial domain. Finally, subsection C.6 will show how the spatial scale of tuning correlation as a function of distance is quantitatively determined by the spatial scale of 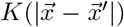 and 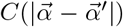 jointly (as in Fig. 4).

#### C.1 Circuit and activity assumptions required for asympotic analysis

We introduce variable 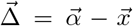 as the location of the receptive field in the LGN relative to the location of the cortical neuron at position 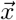. To proceed we perform spatial Fourier transforms of the eigenfunction problem in Eq. (15) with respect to both cortical coordinate 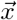 and receptive field coordinate 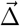, with corresponding variables in Fourier domain to be 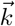 and 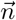, respectively (i.e., 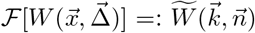). Motivated by numerical simulations of the network, we restrict ourselves to the solution of 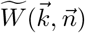 with a single characteristic spatial frequency 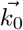 over cortical space 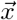, allowing us the simplification:

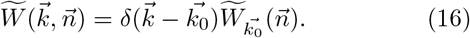

In this case Eq. (15) becomes (See Appendix B for details):

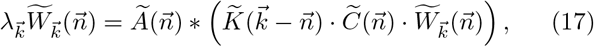

where ∗ denotes convolution, the subscript 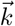 indicates the characteristic spatial frequency, and 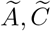, and 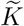 represent the Fourier transforms of functions 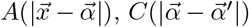 and 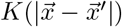, respectively. In the following analysis we denote the polar coordinate of two-dimensional variable 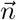 in Fourier domain as (*ρ, ϕ*), and the maximum radial mode of 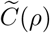 and 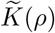 to be 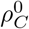 and 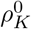, respectively (Fig. 5A, 5B and 5C).

**FIG. 5.**
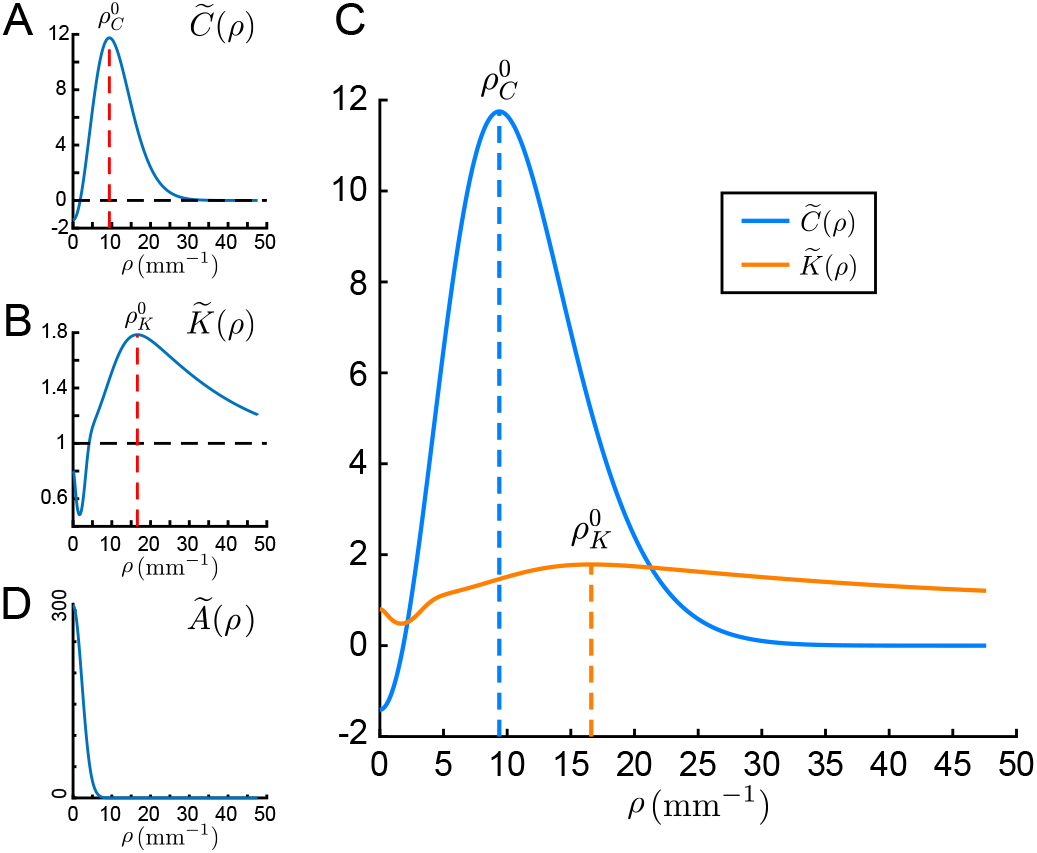
The three main circuit components of the learning equation in the spatial Fourier domain: 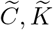 and 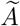. (A) The circular symmetric, band-pass LGN-LGN correlation function 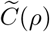. The red dashed line marks the maximal radial mode 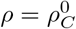. (B) The intracortical recurrent interaction 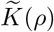. The red dashed line marks its maximal radial mode 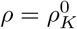. (C) Overlap of panels (A) and (B), where (i) 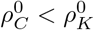 since 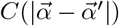 is broader than 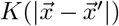 in the spatial domain; (ii) 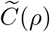 is sharply peaked near 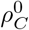 as required by Assumption 1; (iii) 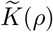 is smooth and flat near 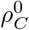 as required by Assumption 2. (D) The low pass arbor function 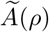, which is sharply peaked near *ρ* = 0 as required by Assumption 3.

The asymptotic regime of mouse V1 that we consider is:

i. The effective intracortical recurrent interaction 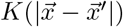 is much narrower than the LGN correlation function 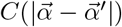 in physical space (Fig. 2C, 2E),
ii. The arbor function 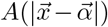 is spatially broad (Fig. 2B),

This regime leads to three asymptotic Assumptions on 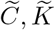 and 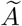 in the Fourier domain (where recall that broad functions in physical space become narrow in the Fourier domain). We mathematically link these three Assumptions through a common small parameter *ϵ >* 0.

Assumption 1

The circularly symmetric, band-pass LGN correlation function 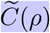 is sharply peaked near its maximal radial mode 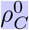. Mathematically, we write this as:

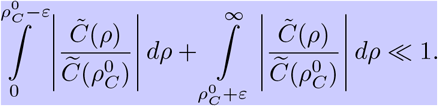

Our standard parameters satisfy this condition (Fig. 5A and 5C), since in general 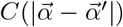 is a broad Mexican-hat-like function in physical space (Fig. 2C).

The intracortical recurrent interaction function 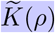 is broader than 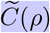, i.e.

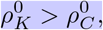

and is smooth and flat near 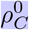, i.e.:

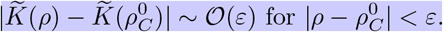

Again this is satisfied in our network (Fig. 5B and 5C, near the cross of dashed blue line and orange curve), since 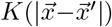 is a narrow Mexican-hat-like function in physical space (Fig. 2E), such that 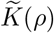 will be broad and with low amplitude (i.e. smooth and flat) in Fourier space.

Assumption 3

The arbor function 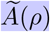 is sharply peaked near *ρ* = 0, i.e.:

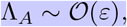

and

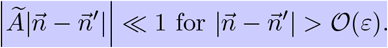

where Λ_*A*_ is the standard deviation of 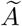:

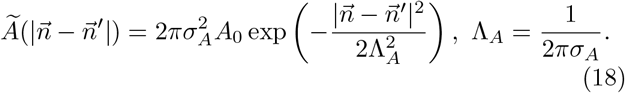

Since *σ*_*A*_ is broad in physical space (Eq. (8)), Fourier transform leads to a small Λ_*A*_ (Fig. 5D).

As we will show, when networks satisfy these three Assumptions, we can focus on the solutions of leading eigenfunctions 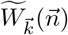 that are ~ *O*(1), and discard *O*(*ε*) and higher order terms (unless *O*(1) terms are trivial constants). We remark that it is possible to assume three small parameters, one for each assumption (i.e *ϵ*_1_, *ϵ*_2_, and *ϵ*_3_). However, in this case the analysis becomes overly cumbersome, while providing limited extra insight.

#### C.2 Separation of radial and angular variables

When Assumption 1 is applied to Eq. (17) the multiplication of a sharply peaked 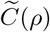 will constrain 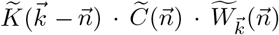 to be also sharply peaked near 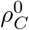. Further, when applying Assumption 3, the convolution of sharply low-pass 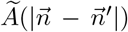 will only slightly blur 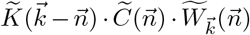 in the Fourier domain. As a result, any leading eigenfunction 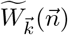 will be sharply peaked near 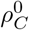, i.e.

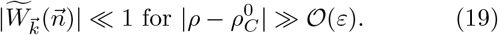

We remark that this is a result of the LGN correlation function *C* and the arbor function *A* being spatially broad.

Assumption 2 gives us the following simplification:

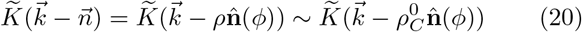

(where the unit vector 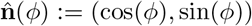), i.e. 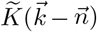 is approximately radially constant at 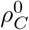, and only depends on the angular coordinate *ϕ*. This is a result of the intracortical recurrent interaction in L4, *K*, being spatially narrow.

Substituting the approximations in Eqs. (19) and (20) into the integral form of Eq. (17) yields:

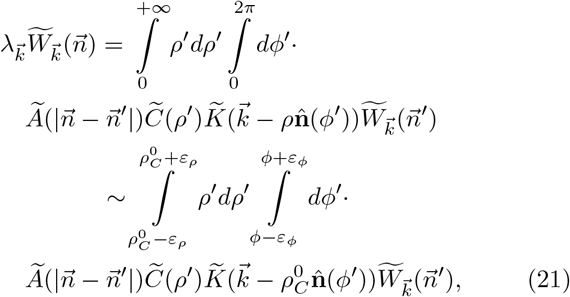

where due to Assumption 1 and Eq. (19) we have that the integration limits of *ρ*^*′*^ are within 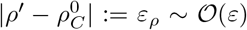. Further, due to Assumption 3, 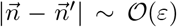, so that |*ρ*−*ρ*^*′*^| ~ *O* (*ε*) and |*ϕ*−*ϕ*^*′*^| := *ε*_*ϕ*_ ~ *O*(*ε*) give the integration limits of *ϕ*^*′*^.

The form of Eq. (21) permits a separation of variables approach to solve Eq. (17). Consider the convolution kernel 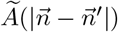 (Eq. (18)) in polar coordinates (*ρ, ϕ*):

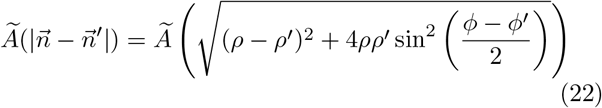

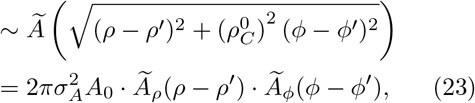

where

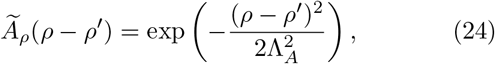

and

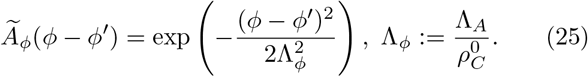

This uses the fact that in Eq. (22) both (*ρ* − *ρ*^*′*^)^2^ and 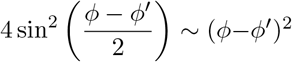 are *O*(*ε*^2^) when both |*ρ*−*ρ*^*′*^| and |*ϕ* −*ϕ*^*′*^| are *O*(*ε*). In total, we then have the approximation 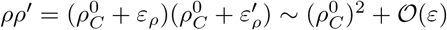.

We take the solutions of 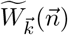 to be separated radially and angularly:

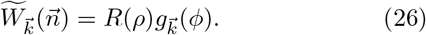

This allows Eq. (21) to be separated into a radial part

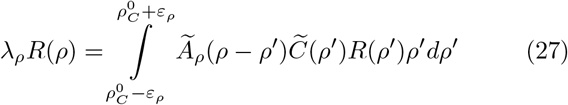

(where *λ*_*ρ*_ is the radial part eigenvalue) and an angular part

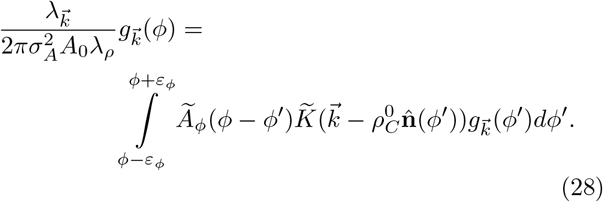

In the next two subsections we study the eigensystem for the radial and angular components separately.

#### C.3 The radial eigenfunction and eigenvalue

In this section we will solve the radial eigenfunction problem in Eq. (27). Assumption 1 allows us to consider the Laplacian approximation to 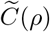 around its peak 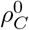:

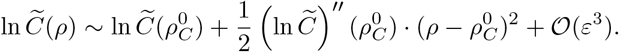

This gives us a Gaussian approximation to 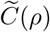 around 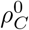 :

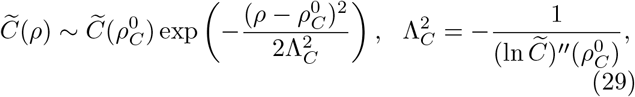

where the expression of 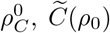 and Λ_*C*_ can be determined by the form of 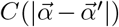 in Eq. (9) (see Eqs. (66), (67) and (68) in Appendix C for details).

Since 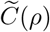 is a one-dimensional Gaussian function centered at 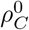, and the convolution kernel 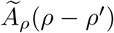 is also a one-dimensional Gaussian function, then the solution of *R*(*ρ*) is also a one-dimensional Gaussian function centered at 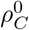:

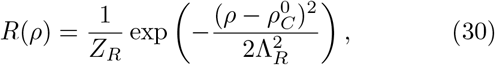

where *Z*_*R*_ is a normalization factor.

Substituting Eq. (30) into Eq. (27) determines the spatial scale of the eigenfunction *R*(*ρ*) to be (see Appendix C for details):

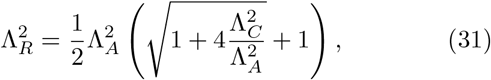

and, finally the radial eigenvalue becomes:

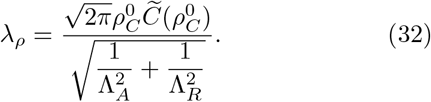

#### C.4 The principal angular eigenfunction and eigenvalue

In this section we will solve the angular eigenfunction problem given by Eq. (28). We denote

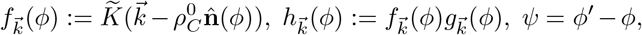

so that Eq. (28) reduces to:

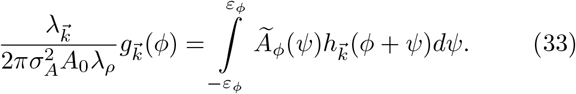

Within the range of |*ψ*| ≤ *ε*_*ϕ*_ ~ *O*(*ε*), we take a Taylor expansion of 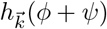:

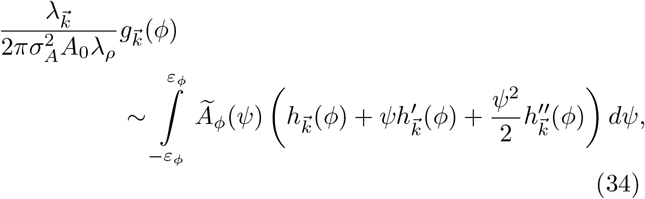

such that the convolution is approximated as differentiation over a small range of *ψ*. Substituting the form of 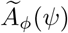 (Eq. (25)) into Eq. (34) yields

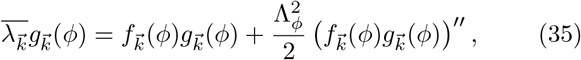

where

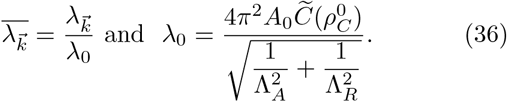

By combining Assumption 3 and Λ_*ϕ*_ ~ *O*(*ε*) we have that the solution of the principal eigenfunctions and eigen-values of Eq. (35) can be well approximated by the solution of quantum harmonic oscillator problem when the quantum number *l* = 0 (see Appendix D for details).

The principal eigenvalue for each characteristic spatial frequency 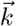 is (Fig. 6):

**FIG. 6.**
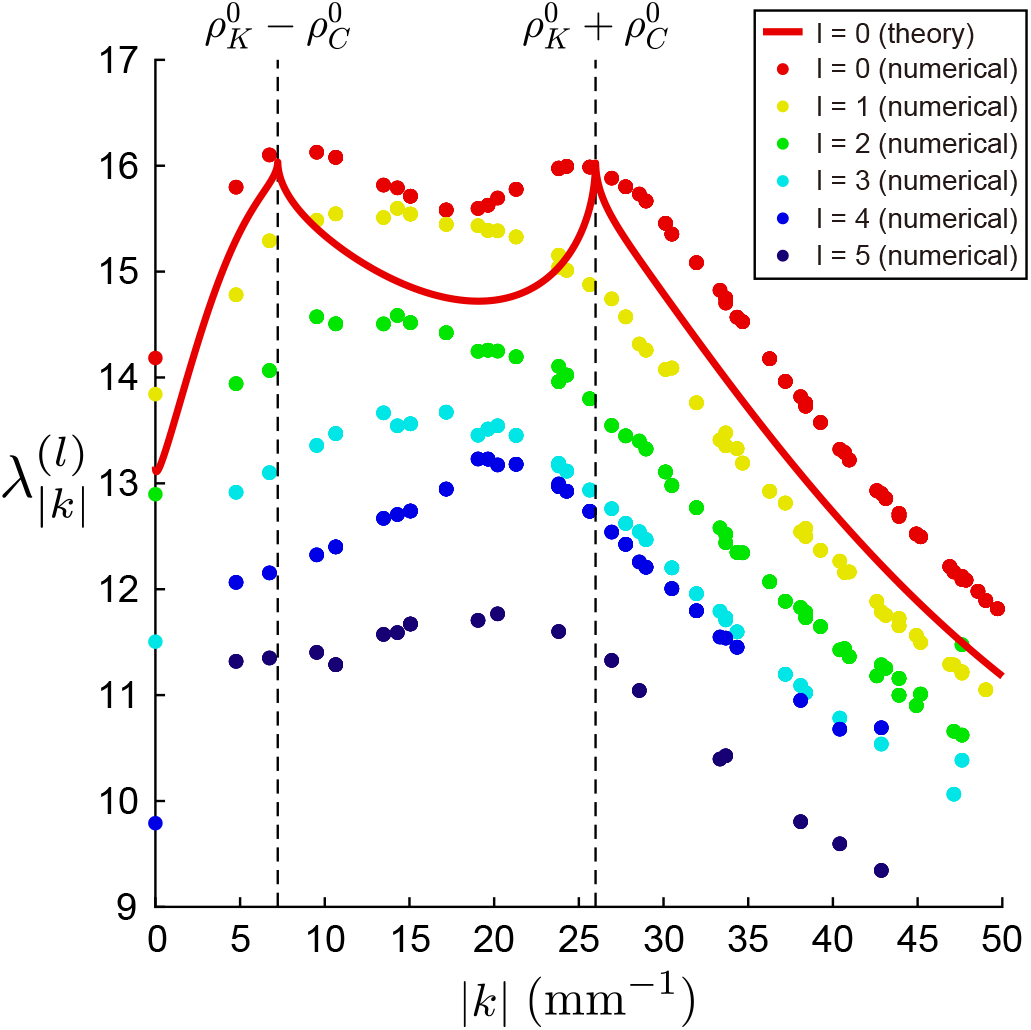
Eigenvalue 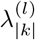 over the modulus of characteristic spatial frequency |*k*| of the corresponding eigenfunction with quantum number *l*, from quantum harmonic oscillator based theory (when *l* = 0, in red curve) and numerical solutions (colored dots over different *l*).

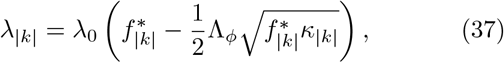

where |*k*| and *ϕ*_*k*_ are the modulus and argument of 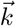. Here we have introduced the notation:

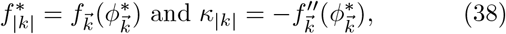

with 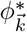 to be a local maxima of 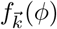 (See Appendix D for details, in particular for the form of the corresponding eigenfunctions 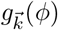 given in Eq. (100)).

A core prediction of our theory is that the most principal eigenvalue *λ*_|*k*|_ (Eq. (37)) attains (locally) maximum values at:

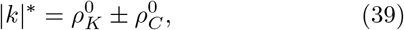

(see Appendix D for details). In other words, the principal eigenfunction that corresponds to the maximal eigenvalue has a spatial mode 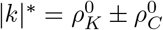. This spatial frequency is jointly determined by the fine spatial scale of intracortical recurrent interaction 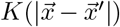 (characterized by 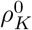) and the broad spatial scale of the LGN correlation function 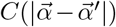 (characterized by 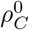). Satisfyingly, this prediction is verified in numerical solutions of principal eigenvalues (Fig. 6, *l* = 0 eigenvalue). We note that the eigenvalues for higher modes (Fig. 6; *l* = 1, 2, 3 …) do not show local maxima at 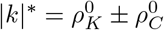. These eigenvalues can only be computed numerically, as we lack theory for their calculation.

In the following subsection we will show how these characteristic spatial frequencies |*k*|^∗^ quantitatively determine the shape and spatial scale of tuning correlation as a function of spatial distance (seen in experiment in Fig. 1 and simulations of the learning model in Fig. 3E).

#### C.5 Principal eigenfunctions in circular harmonic bases

Given the radial and angular parts of the principal eigenfunction in sections C.3 and C.4, we next use the inverse Fourier transform of 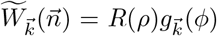 to go from the Fourier domain 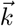 and 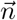 (with polar coordinate (*ρ, ϕ*)) back to spatial domain 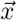 and 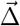 (with polar coordinate (*r, θ*)). The complete form of principal eigenfunctions over 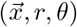 is then (see Appendix E for details):

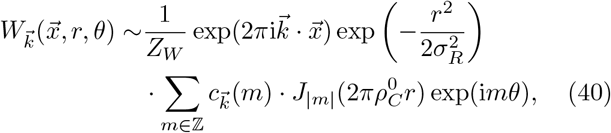

where *Z*_*W*_ is the normalization factor; *J*(·) represents Bessel functions of the first kind (example eigenfunctions shown in Fig. 7D).

**FIG. 7.**
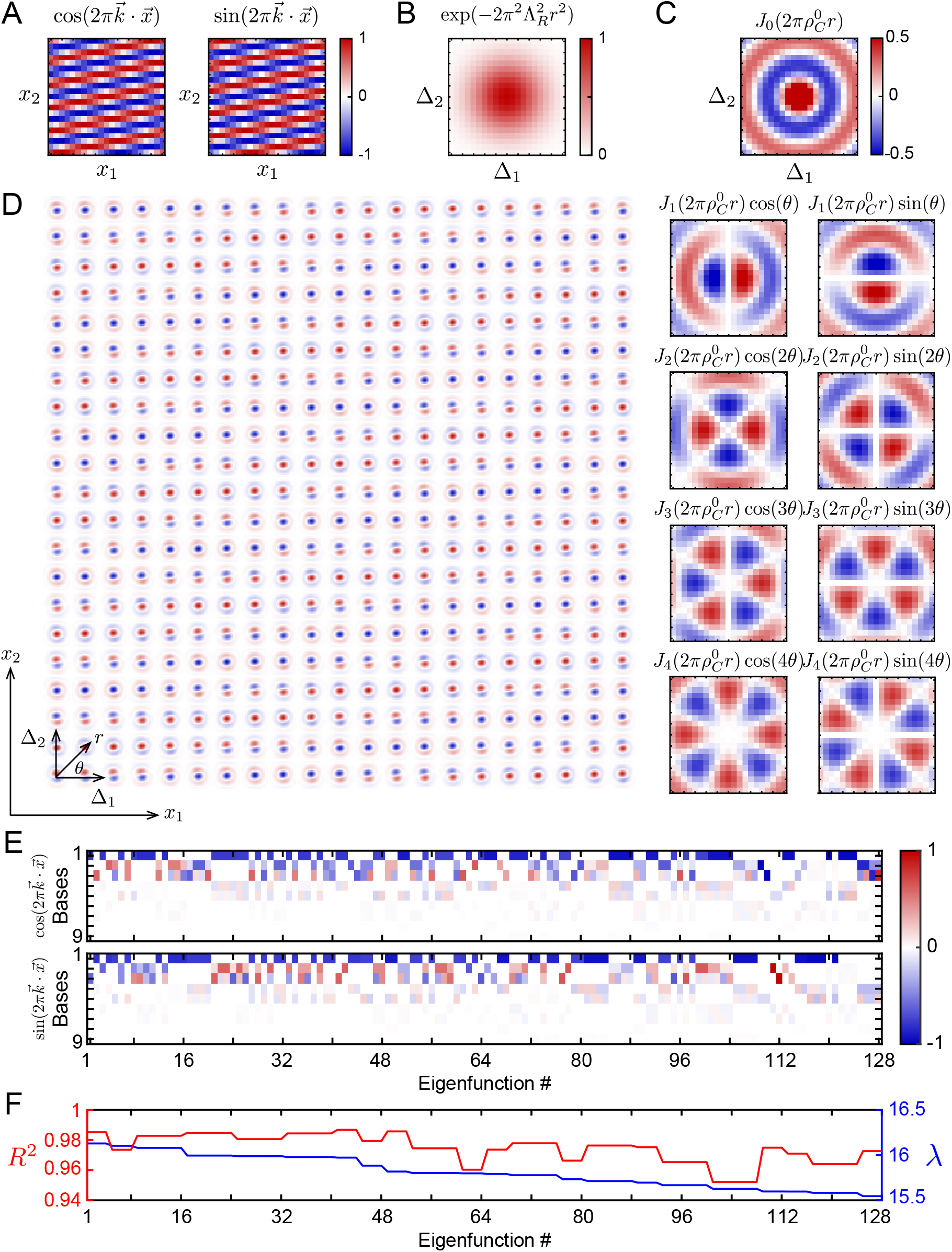
Schematic of principal eigenfunctions in the circular harmonic bases. (A) Schematic of the spatial plane wave term 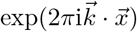 with 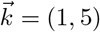 wavenumbers (converted from complex form to real form; also for all following panels). (B) Schematic of the Gaussian envelope function term 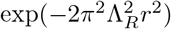. (C) Schematic of the ‘circular harmonic’ bases 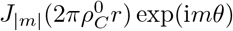 for |*m*| ≤ 4. (D) Example of one principal eigenfunction with maximal eigenvalue, with 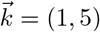 wavenumbers. Note the coordinate 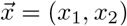 of each small squared panel across cortical neurons, and 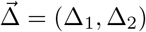 for translated receptive field location of each neuron, as well as its polar coordinate (*r, θ*). (E) The coefficients 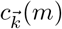 for the 9 real form circular harmonic bases in (C), for the first 128 principal eigenfunctions. (F) Red: The *R*^2^ between analytical form and numerical simulation of first 128 principal eigenfunctions; Blue: The first 128 principal eigenvalues (from numerical simulation).

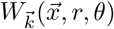 contains three factors:

i. The term 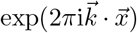 is the spatial plane wave over cortical space with characteristic spatial frequency 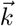 (Fig. 7A).
ii. The term 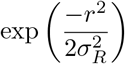 is a Gaussian envelope function on the receptive field (*r, θ*), which is common for all locations 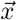 on the cortical space (Fig. 7B). The standard deviation is 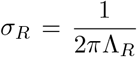 (with Λ_*R*_ as Eq.(31)).
iii. The linear combination of bases 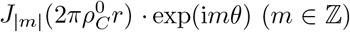, which are the circular harmonic function bases (two-dimensional analog of spherical harmonic bases) of the receptive field of each neuron (Fig. 7C). For the linear combination coefficients 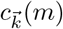 they are associated with the form of principal angular eigenfunctions 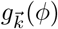 (see Fig. 7E for the coefficients of first 128 principal eigenfunctions with |*m*| ≤ 4; see Eq. (128) in Appendix E for details).

This eigen representation does capture the emergence of Gabor function-like receptive fields (Fig. 7D), as was observed in simulations (Fig. 3A). Note that Eq. (40) can be converted to real functions (as shown in Fig. 7) since 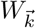 and 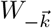 are conjugate pairs. They quantitatively match numerical solutions of principal eigenfunctions well (Fig. 7F).

#### C.6 Determinant of the spatial scale of micro-clustered organization of tuning correlation

In this final subsection we determine the main circuit contributions to the spatial scale of tuning correlations as a function of distance.

Consider receptive field 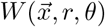 represented as a linear combination of principal eigenfunction bases, with coefficients 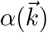 (from Eq. (40)):

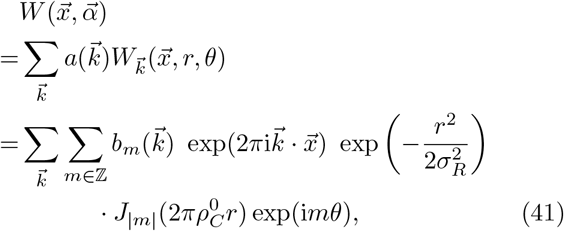

where

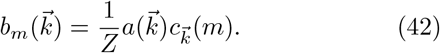

Substitute this representation into the expression for 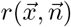 (Eq. (13)), the neuronal response to grating visual stimulus; and define the average of inner product among responses of pair of neurons over specific distance 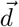:

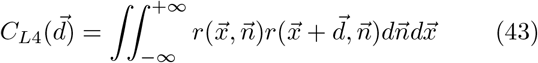

We note that the tuning correlation defined previously is the inner product among whitened neuron responses. We use the subscript ‘L4’ to distinguish 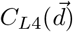 from the LGN correlation 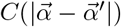.

Finally we have the analytical form of the Fourier transform of 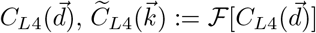: (see Appendix F for details)

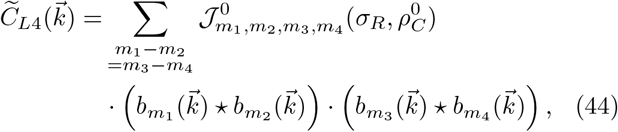

where *m*_1_, *m*_2_, *m*_3_, *m*_4_ ∈ ℤ; ‘⋆’ represents cross-correlation; 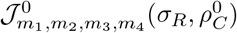 is a scalar coefficient (see Eq. (154) in Appendix F for details).

To further understand how the spatial scale of 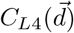 is determined by the characteristic spatial frequency of eigenfunctions with maximal eigenvalue 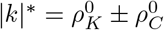 via the inverse Fourier transform of Eq. (44), we make the following parsimonious assumptions:

i. We ignore the differences of functions 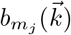 in Eq. (44) across different *m*_*j*_, i.e. we simply consider them to be uniform across different *m*_*j*_ as 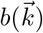. As a result, from Eq. (44) we have the following approximate relationship:

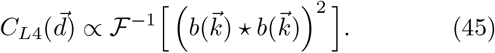
ii. Since Oja’s learning rule selects the principal eigenfunctions with maximal eigenvalue, we assume that the co-efficient 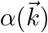 in Eq. (41) and also 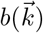 in Eq. (42) will only be non-zero for the characteristic spatial frequency 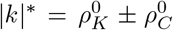 after sufficient learning. In other words, 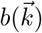 has the following ‘double-ring’ form (Fig. 8A):

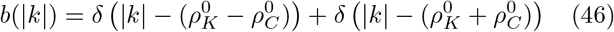

**FIG. 8.**
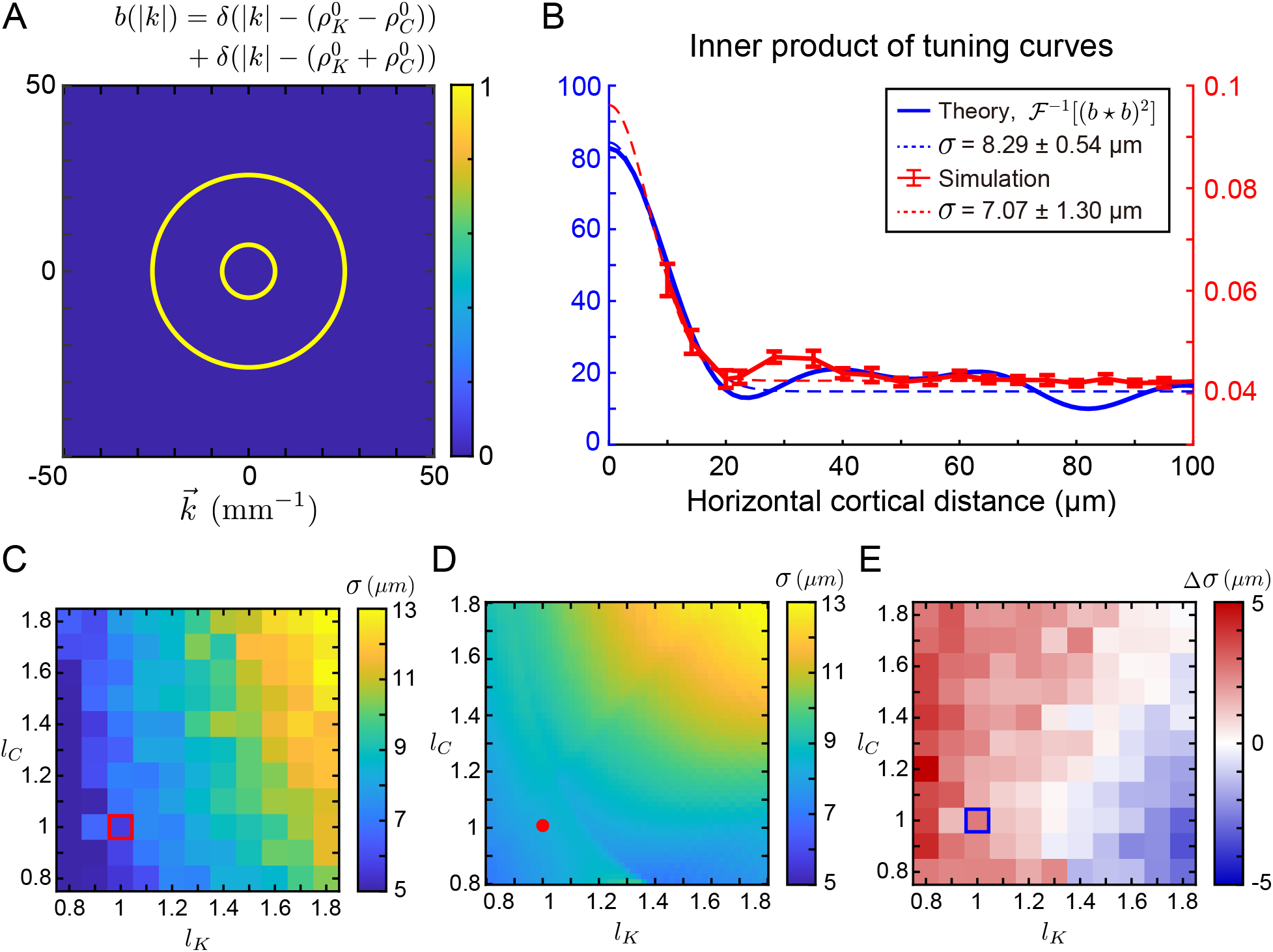
Determinant of the spatial scale of inner product of orientation tuning responses with parsimonious assumptions. (A) The ‘double-ring’ form of coefficient *b*(|*k*|) of eigenfunction bases with different spatial frequency. (B) The inner product of tuning responses as a function of distance between pair of neurons, from numerical simulation (red curve; variant of Figure 3E) and the theory with parsimonious assumptions (blue curve; Eq. (47)). Dashed curves: fitting to Gaussian function; *σ*: the standard deviation / spatial width of fitting (error represents the 95% confidence interval). (C) Same as Figure 4B: the spatial width *σ* of the inner product of tuning responses with different combinations of scaling factors (*l*_*K*_, *l*_*C*_) in numerical simulations. The red box indicates the standard parameters where (*l*_*K*_, *l*_*C*_) = (1, 1). (D) Similar as (C) but generated from theory with parsimonious assumptions. The red dot indicates (*l*_*K*_, *l*_*C*_) = (1, 1). (E) The error Δ*σ* of theoretical *σ* in (D) over numerical *σ* in (C). The blue box indicates (*l*_*K*_, *l*_*C*_) = (1, 1).

Substituting Eq. (46) into Eq. (45), gives the following approximate equation of *C*_*L*4_(|d|) (Fig. 8B; see Appendix F for details):

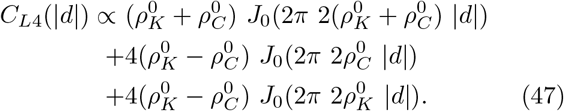

To understand its spatial scale, consider that Bessel function *J*_0_(*z*) ~ 1 − *z*^2^/4 ~ exp(−*z*^2^/4) over small value of *z*. This allows the approximation:

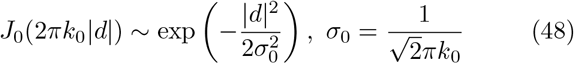

when |*d*| is small, so that *C*(|*d*|) in Eq. (47) contains three components with the following characteristic spatial scales

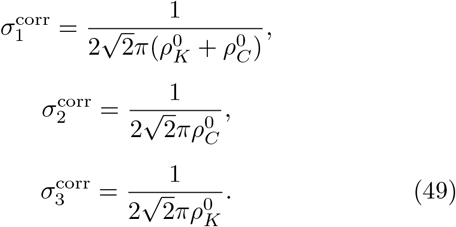

Thus, the spatial scale of intracortical recurrent interaction 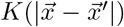 (characterized by 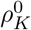) and LGN correlation function 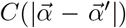 (characterized by 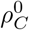) jointly determines the spatial scale of *C*_*L*4_(|*d*|). This is consistent with numerical simulations across various spatial scales of 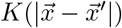 and 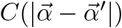 (Fig. 4B, and Fig. 8C ~ 8E). Especially, with standard parameters (Methods; Fig. 2C and 2E; Fig. 5A ~ 5C) of 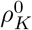 and 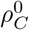, we have that:

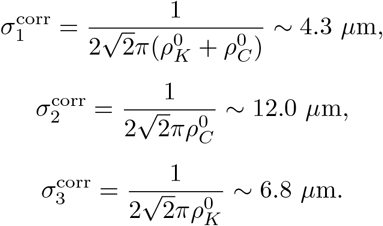

These indicate that the large 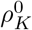 due to spatially confined intracortical recurrent interaction 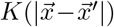 leads to small 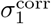 and 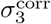, and ultimately the spatially fine *C*_*L*4_(|*d*|), i.e. micro-clustered organization of orientation tuning over fine spatial scale (Fig. 8B).

## III. DISCUSSION

In this work, we investigated a circuit-based mechanism for the development of the fine spatial scale ‘microcluster’ organization of orientation tuning in mouse V1 that we previously identified experimentally ([26]). Building on a framework of synaptic plasticity using Oja’s rule for LGN to L4 connectivity embedded in a spatially structured recurrent L4 network, we combined numerical simulations with analytical theory of the dominant eigenmodes of thalamo-cortical connectivity. Our analysis yields a compact representation of the emergent receptive fields, and provides a mechanistic account of how the spatial scale of the functional map is jointly shaped by correlations of afferent LGN activity and effective short-range intracortical interactions. In particular, the emergence of microclusters requires an effectively micro-scale recurrent interaction, with balanced excitation and inhibition over broad, macro-spatial scales. Together with our previous study on the inheritance of micro-cluster organization from L4 to L2/3 ([26]), these results help to close the theoretical gap by providing an account of how micro-clusters emerge in L4 from thalamo-cortical interactions through developmental processes.

There is a wide variety of theoretical models describing the emergence of cortical functional maps as a self-organization developmental process driven by various learning rules. For orientation tuning preference maps in primary visual cortex, studies have established frameworks of thalamo-cortical wiring plasticity in a spatially structured network where both orientation selective receptive fields and orientation maps are jointly set by (1) the effective intracortical interactions in the cortex; (2) the correlations in afferent LGN activities; and (3) the spatial extent of thalamo-cortical arbor functions [31, 36, 40]. These studies aimed to understand these development processes by pursuing analytical closed-form solutions to the eigen-function problem associated with the linearized equation of learning dynamics (as we have also done). However, these approaches were confined to restrictive limits – taking one of these spatial ingredients to be spatially homogeneous or be asymptotically narrow [41], or assuming all of them are Gaussian functions [36] – which are often far from physiologically plausible conditions in V1.

In our work we address these limitations by showing that the eigenfunction problem becomes analytically tractable in a mouse V1 motivated asymptotic regime for circuit structure. In particular, we introduce a narrow component in circuitry and separate this from the standard broad spatial scales in wiring. First, the intracortical interaction is effectively much more confined over the micro-spatial scale in cortical space than the afferent LGN correlation function, so that the LGN correlation spectrum is sharply peaked around its characteristic mode, whereas the effective intracortical interaction is smooth and flat around this characteristic mode. Second, the thalamo-cortical arbor is sufficiently broad to be well approximated as an effective low-pass filter. Under these two conditions, we obtain an explicit asymptotic solution that yields a compact description of the learned receptive fields in circular-harmonic bases. Importantly, this analysis clarifies the interactions between the model ingredients and the spatial scale of the feature map: the dominant spatial mode, and thus the spatial spread of correlation of tuning responses, is controlled primarily by the LGN correlations and effective intracortical interactions, rather than by the arbor function.

Our theoretical predictions motivate direct experimental validation of the effective recurrent interactions in L4 over fine spatial scales, similar to the validations in L2/3 in our previous study [26]. Measurements such as large-scale cryoEM-based connectomics (for example, the MICrONS project, [42], optogenetic-based perturbation (such as [28]) or high-throughput mapping of synaptic wiring [43], and chronic imaging of synaptic spines in mouse V1 as a direct readout of synaptic plasticity [44], would provide evidence for the regime we analyzed here and further test predicted links between circuitry and micro-cluster functional organization. Beyond circuit measurements, a key implication of our theory is that the correlation structure of visual inputs relayed by LGN could shape the spatial structure of orientation map organization. Testing this prediction will likely require developmental manipulations that alter retinal waves or otherwise decorrelate afferent input activity, as well as controlled perturbations of the animal’s early visual experience.

We emphasize that in the current framework, the intracortical connectivity is treated as a fixed and non-plastic function, and moreover, the excitatory-inhibitory intracortical interactions and ON-OFF LGN units are not explicitly modeled for simplicity. Extending the analysis to allow these factors will be important future steps. In addition, our theory is based on deterministic learning dynamics, which may be disrupted by random synaptic volatility [44]. Understanding the robustness of the proposed mechanism under stochastic plasticity, and its potential relationship to representational drift in V1, will therefore be an important direction for future work [45, 46]. Finally, our analysis provides intuition for how pinwheel-like, micro-clustered, or salt-and-pepper organizations might be selected across species with different spatial scales of input correlations (from LGN or retina) and effective intracortical recurrent interactions. Previous studies have explored related cross-species relationships within a non-developmental theoretical framework [32]. Extending such analyses within a circuit and development based framework may help bridge these perspectives.

## DATA AND COMPUTER CODE AVAILABILITY

All associated codes can be publicly accessed at: https://github.com/brain-math/micro-clusters-L4-learning

## ACKNOWLEDGMENTS

B.D is supported by the NIH grant U19NS107613-01. B.D., NIH grants R01 NS133598 and R01EY037119 and the Simons Foundation Collaboration on the Global Brain. This research benefited from Physics Frontier Center for Living Systems funded by the National Science Foundation (PHY-2317138). Financial support was provided via the National Institute from Mathematics and Theory in Biology (Simons Foundation award MP-TMPS-00005320 and NSF award DMS-2235451).

## METHODS

For the learning model (Fig. 2A), we consider the total number of both L4 neurons and LGN units to be *N* ^2^ = 441, which both uniformly arranged on *N × N* two-dimensional grids where *N* = 21. We take the grid density to be *d* = 0.5 (deg./unit). Given the magnification factor in mouse V1 to be ~ 20 (*µ*m/deg.) [47], this density is equivalent to *d* = 10 (*µ*m*/*unit). Thus, in spatial Fourier domain, spatial frequency 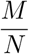 cycles/units is equivalent to 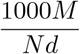 cycles · mm^−1^ (as in Figs. 5, 6 and 8A).

In the learning model given by Eq. (6), we take the learning rate *η* = 1.5 × 10^−4^; the scalar normalization factor *γ*(*t*) is chosen to maintain the Frobenius norm of 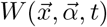 to be *L* = 1 (Eq. (5)).

For the arbor function 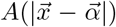 (Eq. (8)), we take *A*_0_ = 1 and *σ*_*A*_ = 3.5 (deg.) [21].

For the LGN correlation function (Eq. (9)), we propose it represents the auto-correlation function of the difference-of-Gaussian shaped function of receptive fields of LGN units (combining both ON-center and OFF-center receptive fields). And we propose the previous measurements of mouse visual contrast sensitivity as a function of the spatial frequency of stimulus [48] reflects the function of LGN receptive field in Fourier domain. Through fitting Eq. (9) to the auto-correlation of the inverse Fourier transform to the measurement of contrast sensitivity as a function of the spatial frequency, we get *C*_1_ = 1.493, *C*_2_ = 0.495; 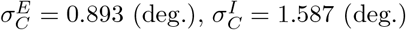 (Fig. 2C).

For the form of the intracortical recurrent connection strength between L4 neurons 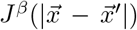 (Eq. (10)) and 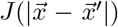 (Eq. (11)), based on hypotheses in previous related studies on the spatial profile of excitatory and inhibitory intracortical recurrent connection strength in mouse V1 [12, 26, 28], we propose 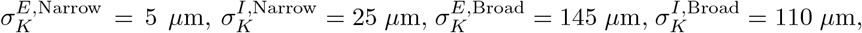, *κ*^*E*^ = *κ*^*I*^ = 0.1, and *A*^*E*^ = 5.05, *A*^*I*^ = − 5.5 (Figure 2E). Notably, the choice *A*^*E*^ and *A*^*I*^ such that *A*^*E*^ +*A*^*I*^ ~ 0 leads to a mostly balanced (canceled) excitation and inhibition over *broad* spatial scales [26].

The global orientation selective index (gOSI, Fig. 3C) is defined by:

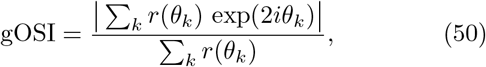

where k is the index among all stimulus orientations, and *r*(*θ*) is the tuning curve responses. *i* is imaginary unit and |·| is modulus. *gOSI* = 0 means no selectivity, and *gOSI* = 1 means perfect selectivity.

For the tuning responses as a function of distance between pair of neurons (Fig. 3E), we calculate the ‘cortical’ distance between neurons on the two-dimensional layer, and the correlation coefficient (i.e. normalized dot product) of tuning curve responses for pairs of neurons.

## APPENDICES Appendix A: Solution of the steady state of linearized L4 dynamics (1)

The steady state of linearized dynamics of L4 neurons in Eq. (1) is:

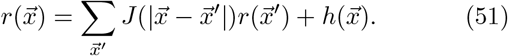

With translational invariant L4 recurrent wiring profile 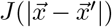 and periodic boundary conditions, the convolutional form Eq. (51) can be solved by Fourier transform:

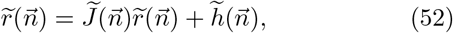

Where 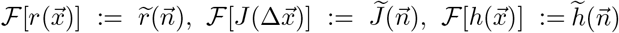. Throughout we take the Fourier transform to be

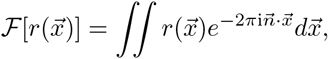

and the inverse Fourier transform to be

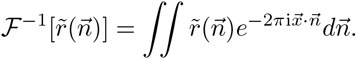

Eq. (52) gives that

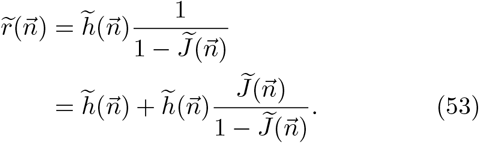

We write the second line since 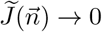 for 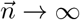, meaning that 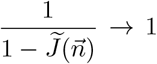 while 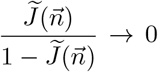 for 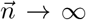. Because of this only 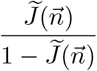 is valid for an inverse Fourier transform. Consider

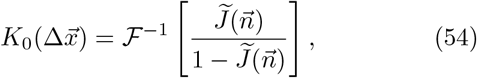

so that the inverse Fourier transform of Eq. (53) is

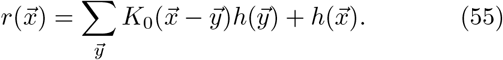

Define the convolution kernel

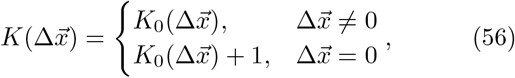

then we have the solution of Eq. (51):

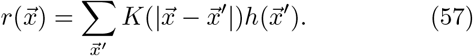

## APPENDIX B Linear operator in the Fourier domain

Consider the original linear (Fredholm) operator in Eq. (15):

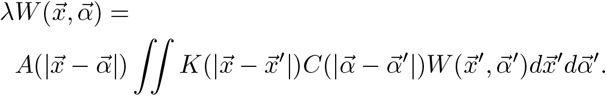

Replace 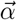 with new coordinate 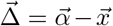, i.e. the location of the receptive field relative to the location of the cortical cell. Similarly, 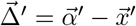. This gives:

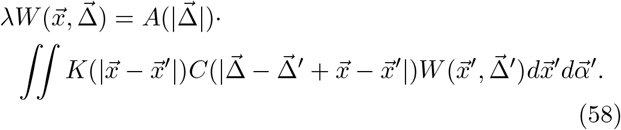

We consider the change of variables 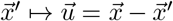 to yield:

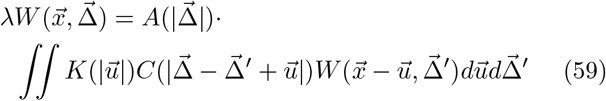

(Note that 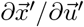 equals to 1 when 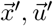 are two-dimensional vectors).

We perform the Fourier transform from 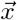 domain to 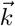 domain, i.e. 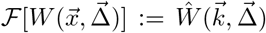. Motivated from numerical simulations of the network, we only care about the solution of 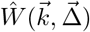 with a single spatial frequency over cortical space 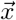:

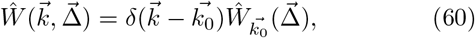

so we have

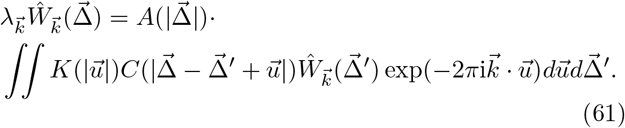

We define 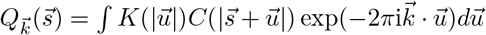, to give:

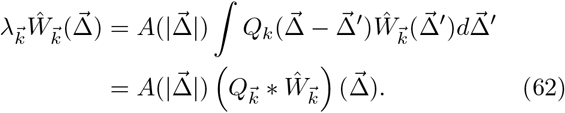

Next we perform the Fourier transform from 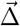 to 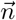 domain, i.e. 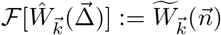 to give:

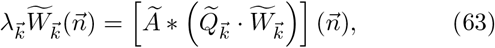

where

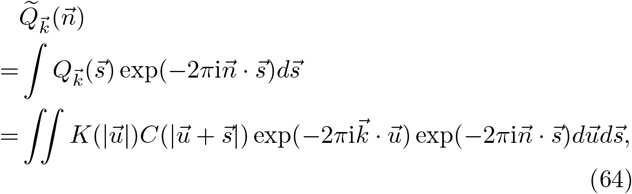

and use the change of variables 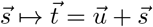 to yield:

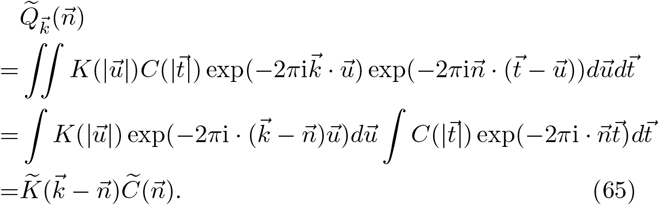

So finally we have Eq. (17):

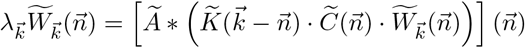

## Appendix C Solution of the radial eigenfunction *R*(*ρ*)

We start from the Laplacian approximation to 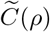 around its peak 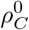 as a Gaussian function as in the main text:

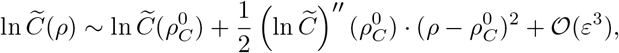

i.e.

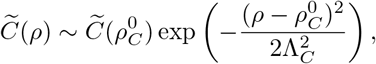

with

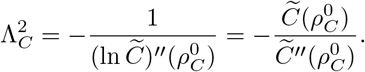

Given the form of 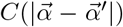 in Eq. (9) we have the location and value of the peak:

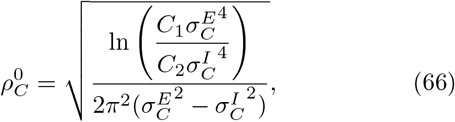

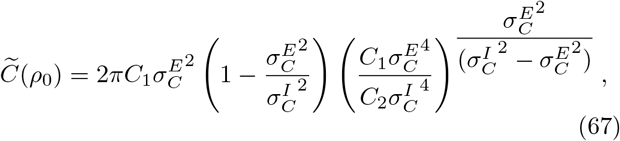

as well as

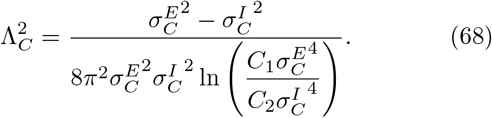

Substitute the form of 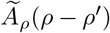 in Eq. (25), the form of 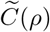 in Eq. (29) and the form of *R*(*ρ*) in Eq. (30) to the right hand side of Eq. (27), we have

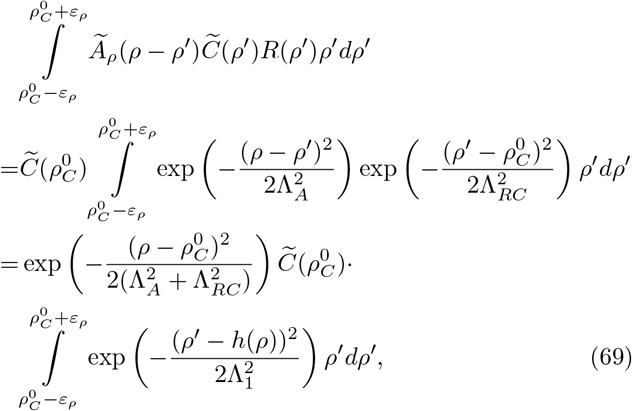

where

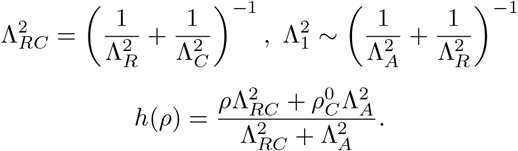

The term

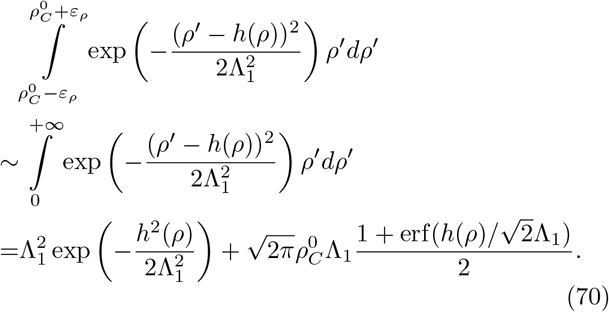

Combining Assumption 3 where Λ_*A*_ ~ *O*(*ε*) and Eq. (19) where Λ_*R*_ ~ *O* (*ε*), we have that Λ_1_ ~ *O*(*ε*). Thus, 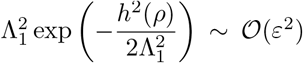 can be discarded, and 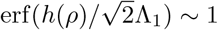.

Thus, Eq. (27) becomes

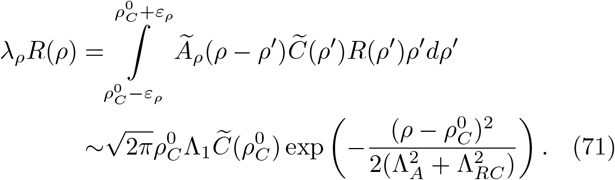

which requires

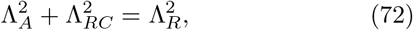

so we have solution of Λ_*R*_ as in Eq. (31):

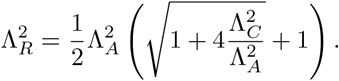

and solution of *λ*_*ρ*_ as in Eq. (32):

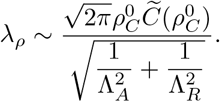

## Appendix D Solution of the eigenvalue and corresponding principal angular eigenfunction 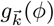

For convenience we repeat Eq. (35):

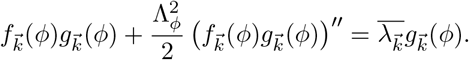

Since Λ_*ϕ*_ ~ *O*(*ε*), when 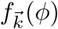 is small nearby its local minima, 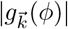 has to be small too (otherwise the absolute value of left hand side will be small but large for right hand side); and when 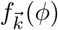 is large nearby its local minima, 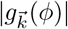 has to be large too (otherwise the absolute value of left side will be large but small for right hand side). Thus, any non-zero part of 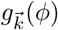 must be around the local maxima of 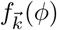.

### Appendix D.1: Local maxima of 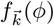

Consider the shape and local maxima of 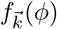 for different spatial frequency 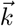:

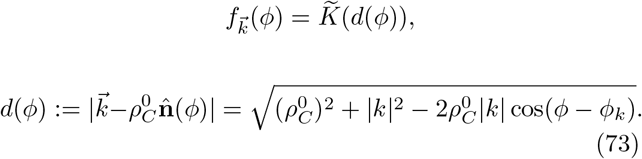

where

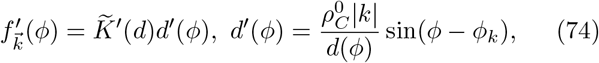

and

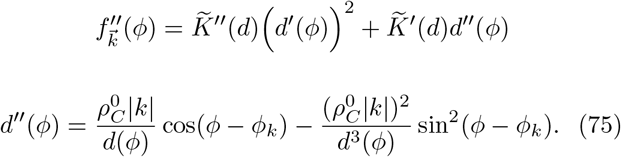

Since 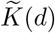 has a local maximum at 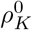 (Figure 5B, 5C), we have that 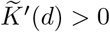 when 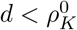, and 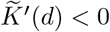 when 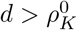.

There are three possible solutions 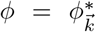 so that 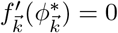:

a. 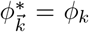 so that 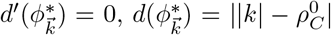, and 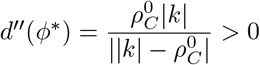. For a local maximum at 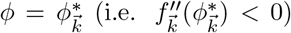 we need 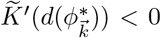, i.e. 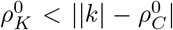. Since 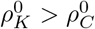, this is equivalent to 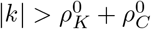.
b. 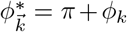 so that 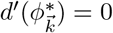, and 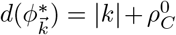, and 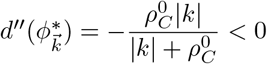. For local maximum at 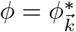 we need 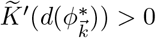, i.e. 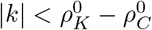.
c. 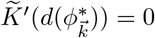, i.e. 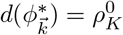. Under the condition that 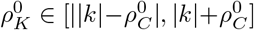, i.e. 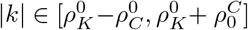, we have that

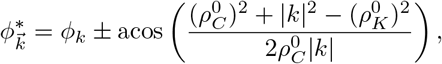

and 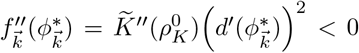 since 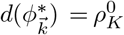 is local maxima and 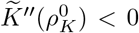. This implies that the two extrema are local maxima.

In summary, with notation of

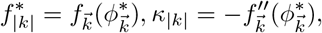

i. When 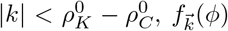 has single, global maximum:

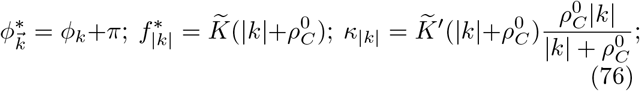
ii. When 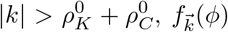 has single, global maximum:

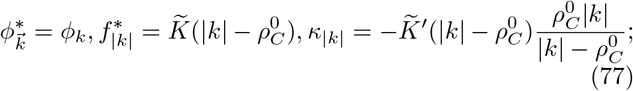
iii. When 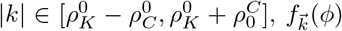 has two local maxima with the same maximal value:

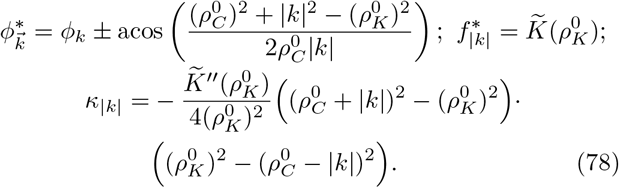

### Appendix D.2: Solution when 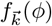 has single local maximum (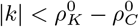 or 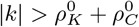)

We first consider the case 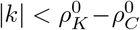 or 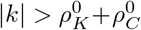 when 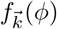 has a single maximum. Consider the ansatz that the solution 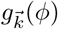 of Eq. (35) is confined nearby the local maxima 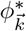, i.e. 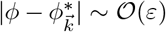, such that 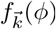 can be well approximated by:

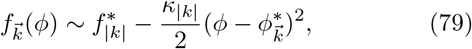

by discarding *O*(*ε*^3^) terms. Rewrite Eq. (35):

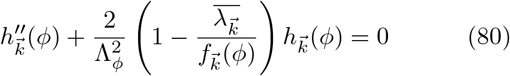

where 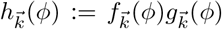. Using Eq. (79) we have the following reduction:

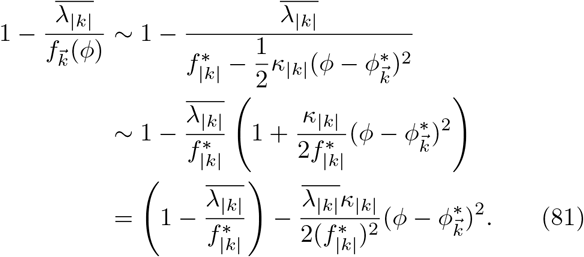

We define

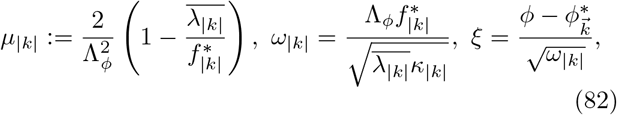

so that Eq. (80) becomes:

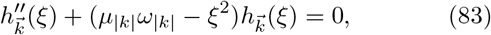

which is equivalent to the quantum harmonic oscillator ([49]). The solutions are:

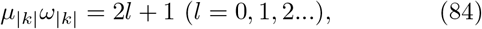

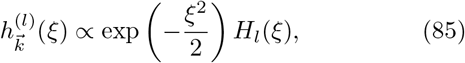

where *H*_*l*_(*ξ*) is Hermite polynomial. Substituting Eq. (82) into Eq. (84) we have:

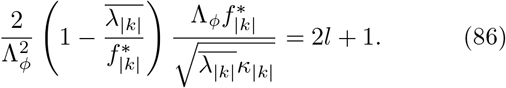

This yields the relation:

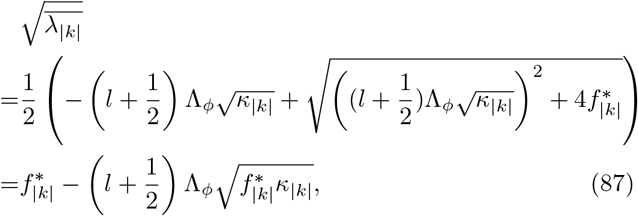

when 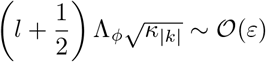.

In summary, the eigenvalues and angular eigenfunctions of Eq. (35) are

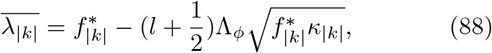

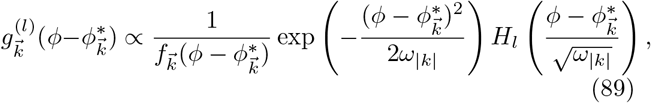

where

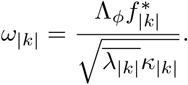

Note that *ω*_|*k*|_ ~ *O*(*ε*) (due to Assumption 3 and Λ_*ϕ*_ ~ *O*(*ε*)). Thus, when the quantum number *l* = 0 we have:

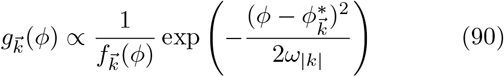

is confined to the region of 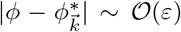 so that the ansatz of Eq. (79) holds. While for *l* ≥ 1, 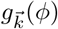 is the product of Gaussian function and a higher order Hermite polynomial, and can thus be away from 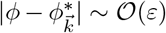 so that the ansatz of Eq. (79) fails.

### Appendix D.3: Solution when 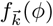 has two local maxima ( 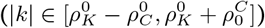) when *l* = 0

In this subsection we ignore subscripts 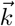 or |*k*| for simplicity. Consider the case 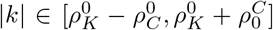 when *f*(*ϕ*) has two local maxima (one to the left of *ϕ*_*k*_, and one to the right of *ϕ*_*k*_):

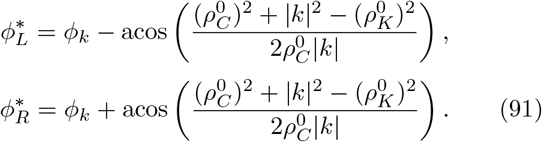

When 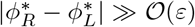, consider the functions *g*^*L*^(*ϕ*) and *g*^*R*^(*ϕ*) in the form of Eq. (89) centered at 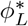 and 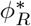:

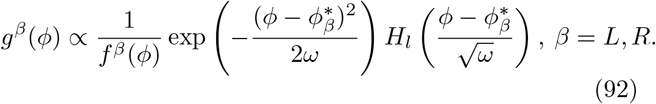

Here we have introduced the notation:

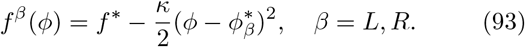

We note that *f* ^*L*^(*ϕ*) and *f* ^*R*^(*ϕ*) share the same *f* ^∗^ and *κ* as in Eq. (78), so that *g*^*L*^(*ϕ*) and *g*^*R*^(*ϕ*) will satisfy

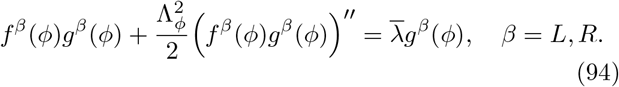

We claim that *g*(*ϕ*) = *g*^*L*^(*ϕ*) + *g*^*R*^(*ϕ*) is an approximate solution of Eq. (35) with bimodal *f*(*ϕ*) and the same eigen-value 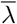:

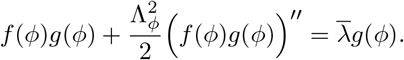

For this to be valid we require the error

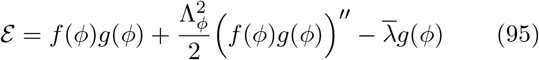

be small. Inserting *g*(*ϕ*) = *g*^*L*^(*ϕ*) + *g*^*R*^(*ϕ*) into the expression for *ℰ* gives:

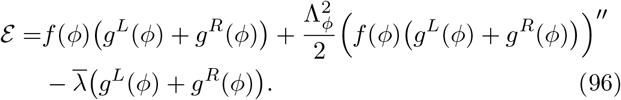

Substituting Eq. (94) into Eq. (96) yields:

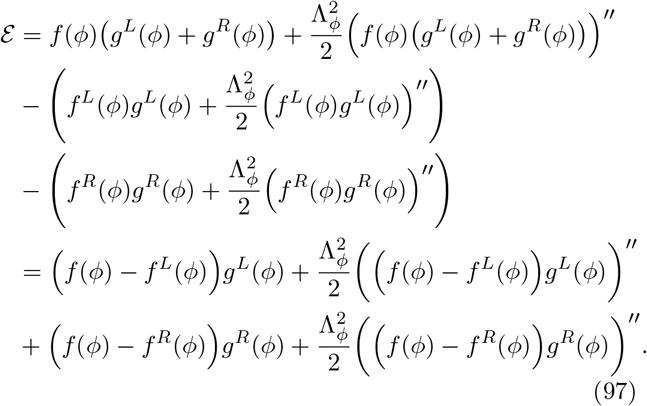

Similar to the case of unimodal *f*(*ϕ*), *ω* ~ *O*(*ε*) so that when *l* = 0, *g*^*β*^(*ϕ*) in Eq. (92) is confined to the region of 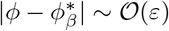 so that

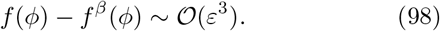

Together with 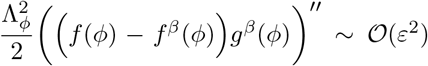 (since Λ_*ϕ*_ ~ *O*(*ε*)), finally we have small ℰ~ *O*(*ε*^2^). In other words, for *l* = 0 we have that *g*(*ϕ*) = *g*^*L*^(*ϕ*) + *g*^*R*^(*ϕ*) is an approximate solution. By contrast, when *l* ≥ 1, *g*^*β*^(*ϕ*) will be no longer confined around 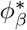, so that |*ℰ*| ≫ 0, then *g*(*ϕ*) = *g*^*L*^(*ϕ*)+*g*^*R*^(*ϕ*) is no longer an approximate solution.

### Appendix D.4: Summary of the solution of eigenvalue and angular eigenfunction

In summary, limited to quantum number *l* = 0 (i.e. principal eigenfunctions and eigenvalues), we have eigen-values (as Eq. (88)):

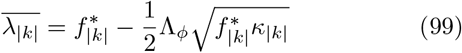

that apply to all ranges of |*k*|. The corresponding eigen-functions are:

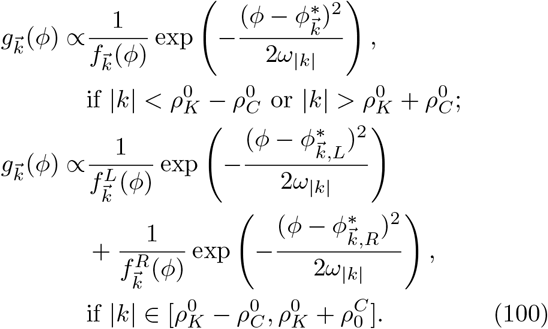

with parameters:

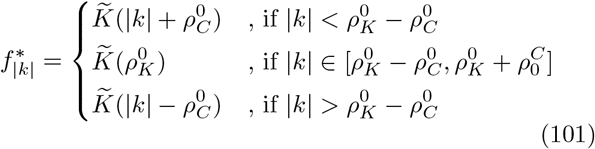

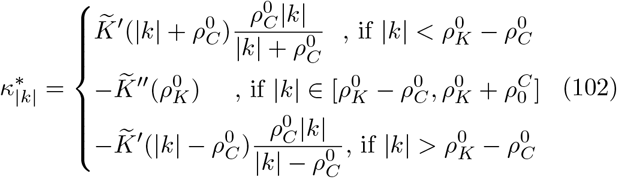

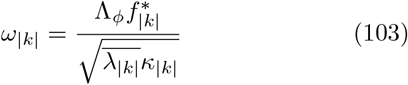

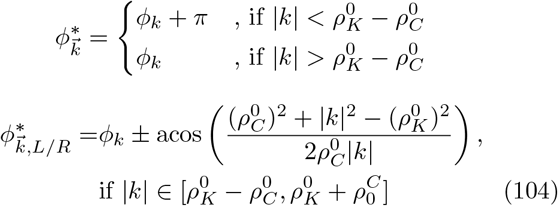

and the asymptotic quadratic approximation of 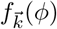 is:

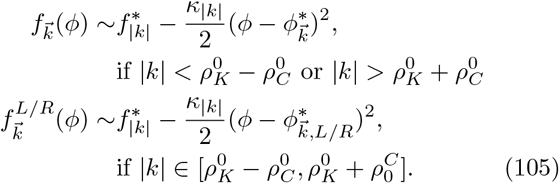

A special case is when 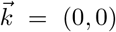, so that 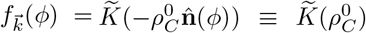 is a constant. Given *g*(*ϕ*) = *g*(*ϕ* + 2*π*), then the solutions are:

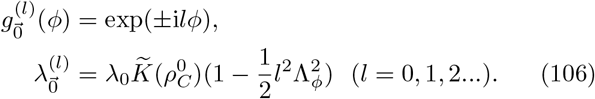

### Appendix D.5: Local maxima of 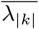 over |*k*|

Finally, we compute the relationship between 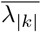 and |*k*|.

First, in Eq. (99) the term 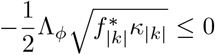, which equals to 0 when *κ*_|*k*|_ = 0 at 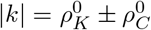, based on Eqs. (76), (77) and (78). For the term 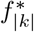 in Eq. (99) we have the following,

a. When 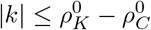,

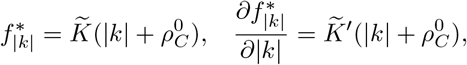

and 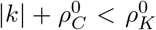. Since 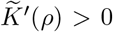 when 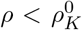, we have 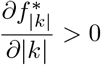. Together with 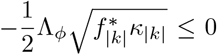, we have 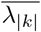 attains maximum 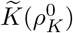 at 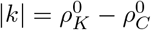.
b. When 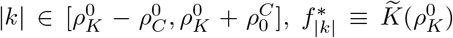, so 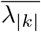 attains maximum 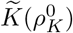 at 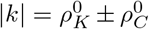 when *κ*_|*k*|_ = 0.
c. When 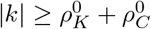,

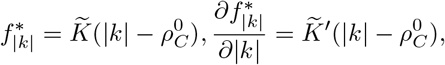

and 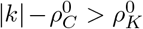. Since 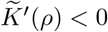 when 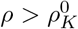, we have 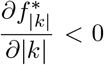. Together with 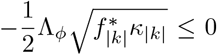. we have 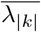 attains maximum 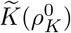 at 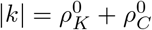.

Thus, for *l* = 0, the maxima of 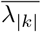 is 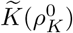, at 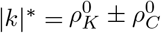 (Fig. 6).

## Appendix E Inverse Fourier transform of principal eigenfunctions

Now that we have the radial and angular parts of principal eigenfunctions 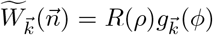, we next consider its inverse transform from Fourier domain 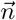 (with polar coordinate (*ρ, ϕ*)) back to spatial domain 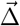 coordinate (*r, θ*)):

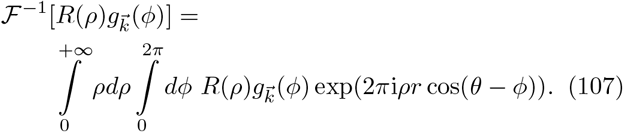

Insert the Jacobi-Anger expansion:

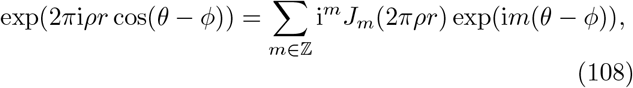

into the expression for 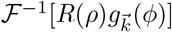 to give:

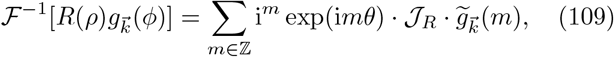

where

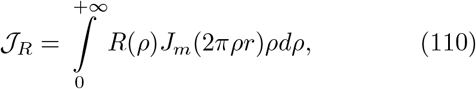

and

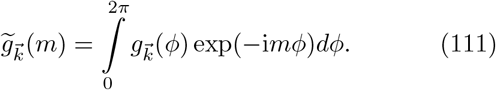

In the expression for *J*_*R*_ we let 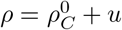 to give:

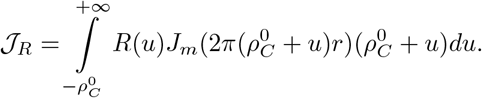

Based on Assumption 1 and that *R*(*ρ*) will be sharply peaked near 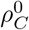 (Eq. (19)), we have that *u* ~ *O*(*ε*). Thus, we consider *O*(1) approximation of *J*_*R*_:

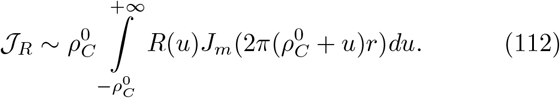

The large-argument asymptotic of Bessel function *J*_*m*_(2*πρr*) (when *r* is large and *ρr* ≫ 1) is:

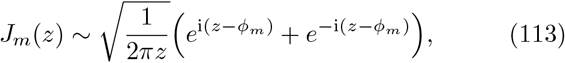

where 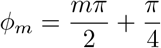. Thus,

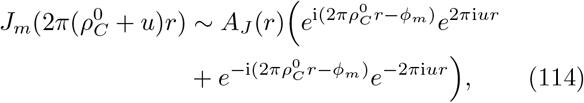

where

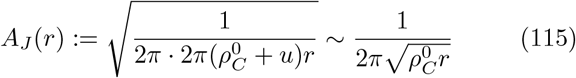

for the *O*(1) approximation. Thus,

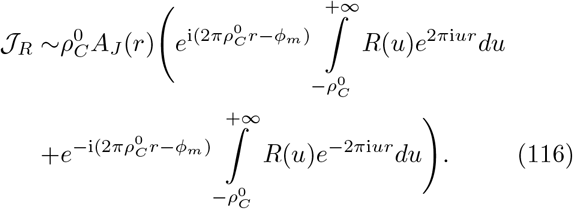

Since *R*(*u*) is sharply peaked, i.e. with standard deviation Λ_*R*_ ~ *O* (*ε*) (due to Assumption 3 and Eq. (31)) that 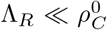, and since *R*(*u*) is an even function (zero-mean Gaussian function), we have that:

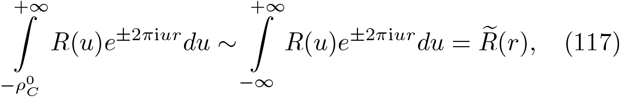

i.e. the Fourier transform of *R*(*u*). Thus,

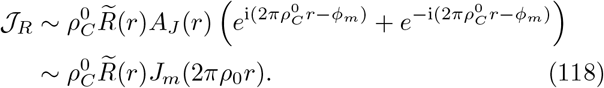

Given *R*(*ρ*) by Eq. (30):

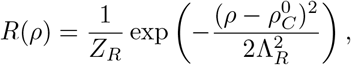

where

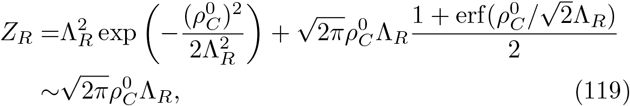

when 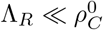. Thus,

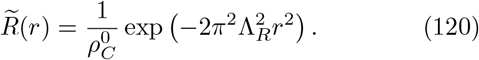

So finally

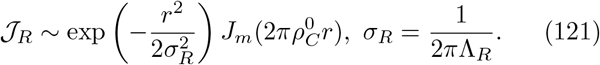

Substituting Eq. (121) into Eq. (109) gives:

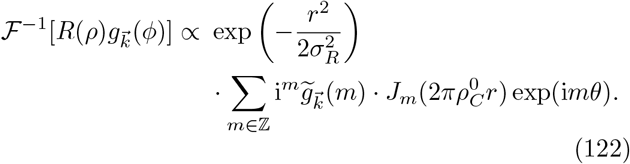

When *m <* 0,

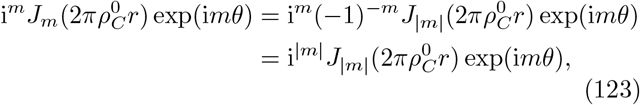

so we have

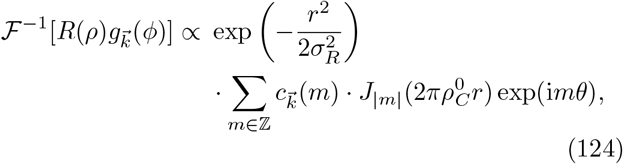

where

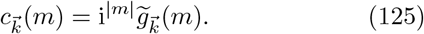

Finally, we are in a position to consider the inverse Fourier transform of Eq. (124) from Fourier domain 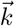 back to spatial domain 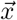 toward the complete form of principal eigenfunction. Given Eq. (16) that each eigenfunction contains only a single spatial frequency 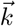, we can simply multiply the factor 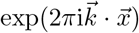 to get the complete form of principal eigenfunction over 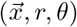 as in Eq. (40):

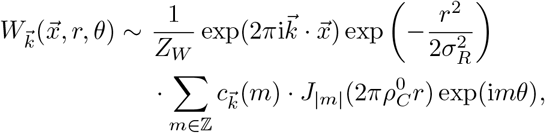

where *Z*_*W*_ is the normalization factor.

The coefficient 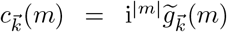 from 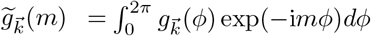, and given the form of 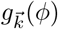 in Eq. (100) we have:

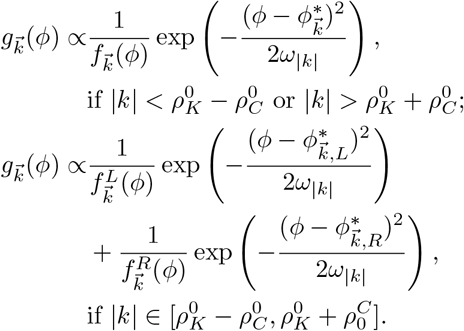

and

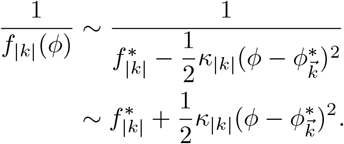

Thus, when 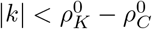 or 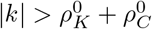 we have:

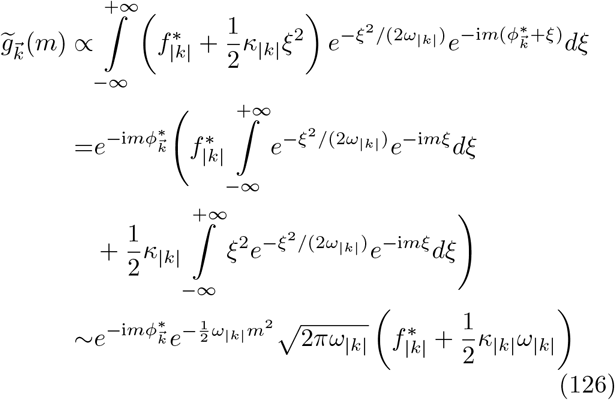

where 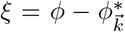, and since *ω*_|*k*|_ ~ *O*(*ε*), the integration over [0, 2*π*] can be extended to (−∞, +∞). When 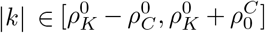, we proceed similar as above to obtain:

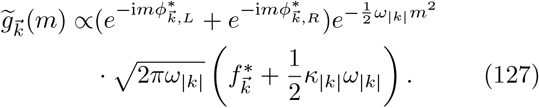

So finally there we have:

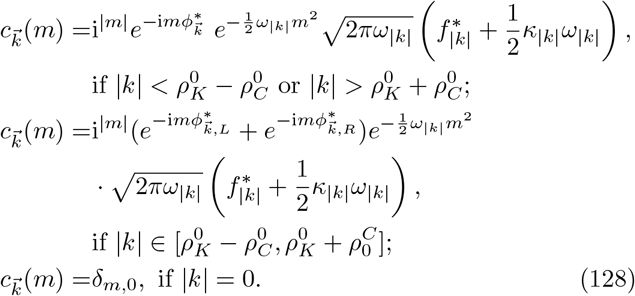

For the special case when 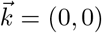 and *l* = 0, since 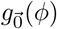 become constant (due to Eq. (106)), the 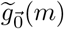 in Eq. (111) is constant when *m* = 0, and 0 when *m ≠* 0. As a result,

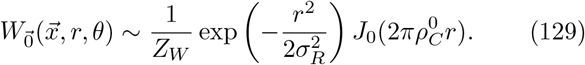

## Appendix F Correlation of orientation tuning responses over space

Consider the receptive field 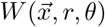 represented in the subspace of principal eigenfunctions (as Eq. (40)):

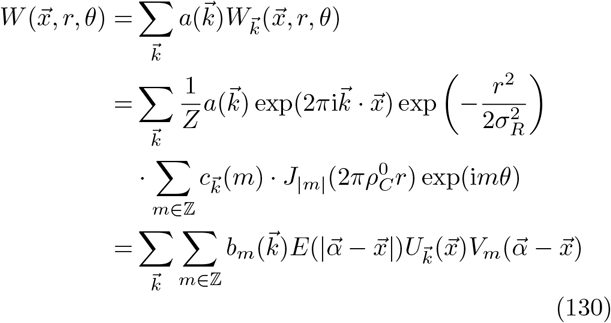

where the coefficient 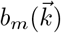 is given by:

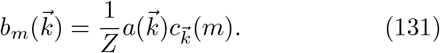

The envelope function is simply:

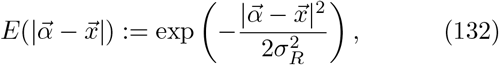

and the circular harmonic basis is given by:

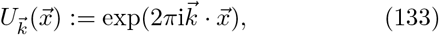

and

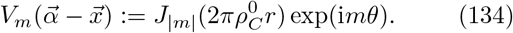

The receptive field 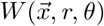, as well as the grating stimuli (Eq. (12))

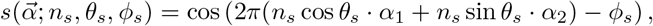

jointly determine the response of L4 neurons (Eq. (13))

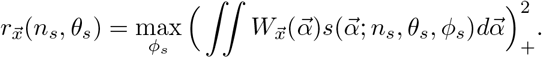

We have the following:

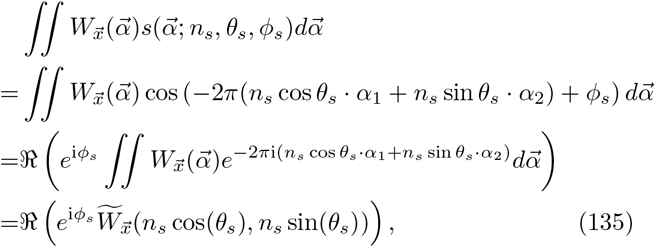

where 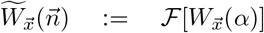. Thus, for 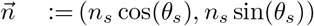 the firing rate obeys:

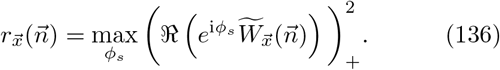

Note that ℜ (*e*^i*ϕ*^*z*) attains maximum value |*z*| when *ϕ* = − arg(*z*). Thus,

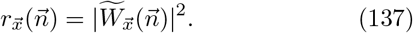

The inner product among responses of neuron at 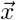 and 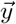: (note that the correlation coefficient of tuning responses defined in the main text is the inner product among whitened neuron responses) is defined as:

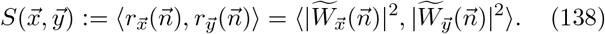

Given that Fourier transform preserves the inner product, and using the Wiener-Khinchin theorem we have:

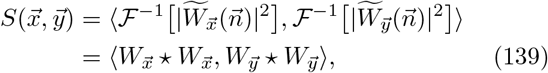

where ⋆ represents cross-correlation. This is why we applied square for neuron responses in Eq. (13), since 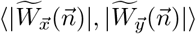 is hard to calculate.

Substituting the expression of 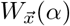 in Eq. (130), we have

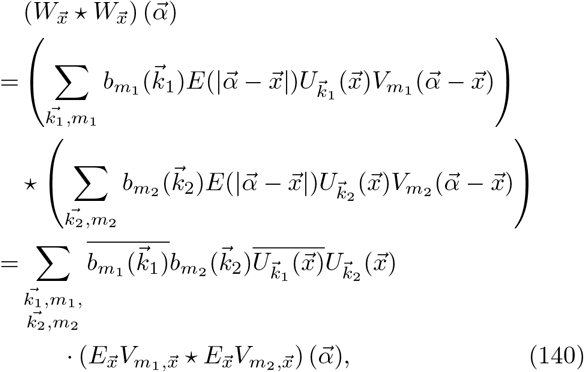

and

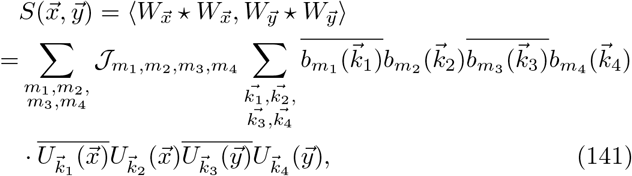

where

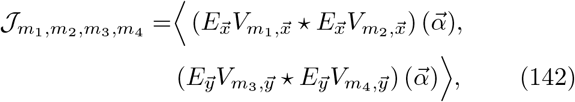

which later we will show to be a constant independent from location 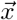 or 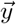.

The average of inner product (Eq. (141)) over certain distance 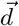 between neuron pair is:

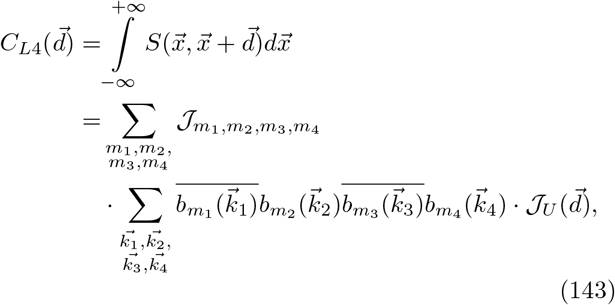

where

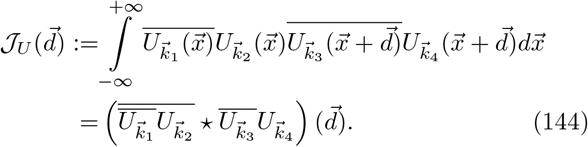

We next consider the Fourier transform 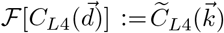. Using the Wiener-Khinchin theorem we have:

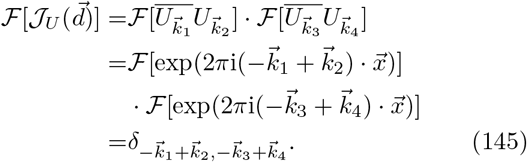

So finally we have:

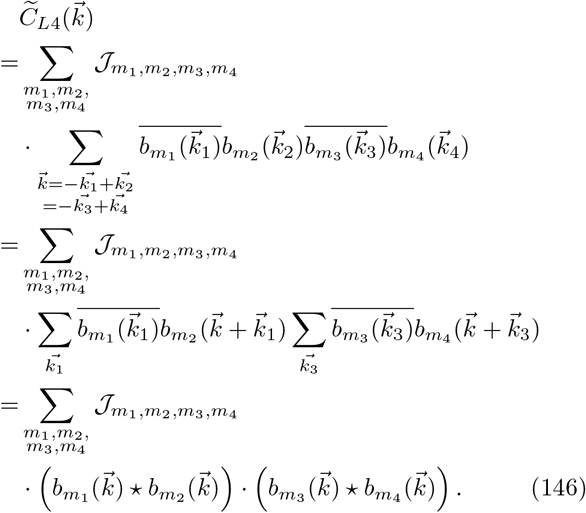

For the term 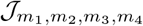 in Eq. (142),

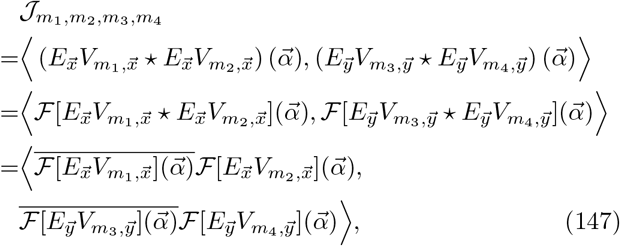

where

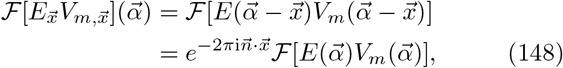

then

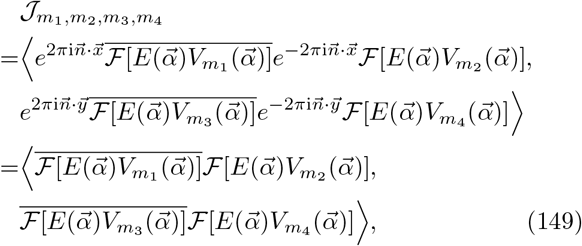

which is a constant independent from location 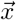 or 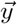.

Consider the Fourier transform from 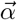 (with the polar coordinate (*r, θ*)) in the space domain to 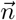 (with the polar coordinate (*ρ, ϕ*)) in the Fourier domain:

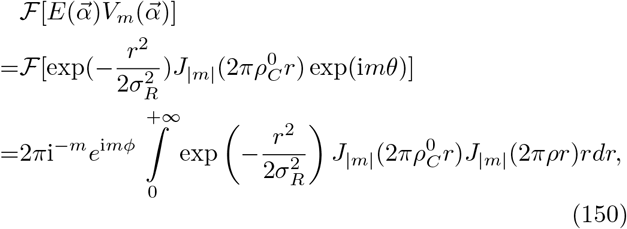

where we have the Weber-Sonine integral ([50]; [51])

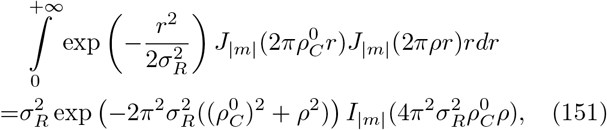

where *I*_|*m*|_ represents the modified Bessel function of the first kind. Thus,

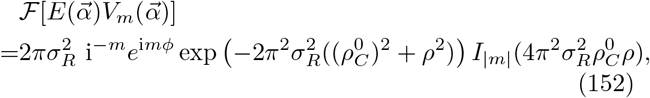

and

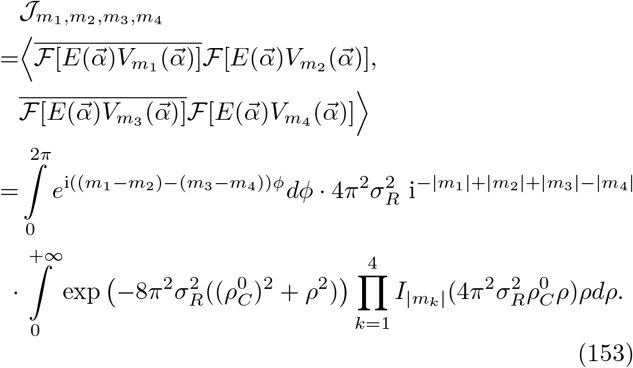

Note that

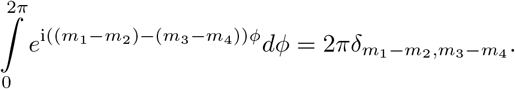

Substituting this expression into Eq. (146) we obtain Eq. (44):

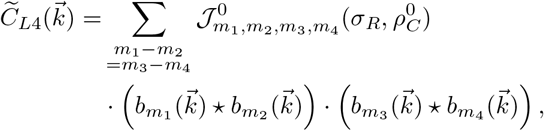

where

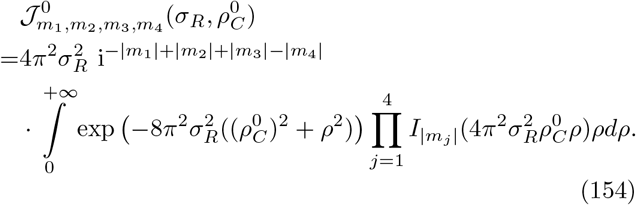

We consider the parsimonious model of 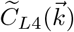, with an oversimplification of the functions 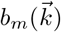 in Eq. (44) to 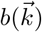 that are uniform across different *m*. We take the following form (as Eq. (46)):

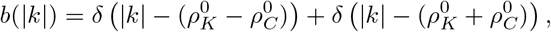

so that the inner product of neural response over spatial distance will be (as Eq. (45)):

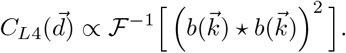

Denote

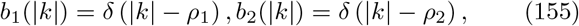

where 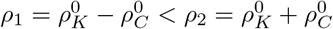, and

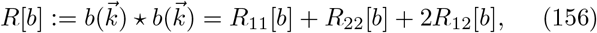

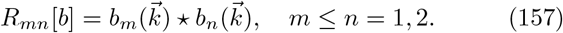

We have

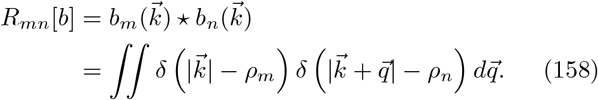

Denote 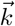 in polar coordinate as (*ρ, ϕ*_*k*_), and 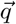 in polar coordinate as (*q, ϕ*_*q*_), then we have

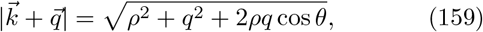

where *θ* = *ϕ*_*k*_ − *ϕ*_*q*_. So,

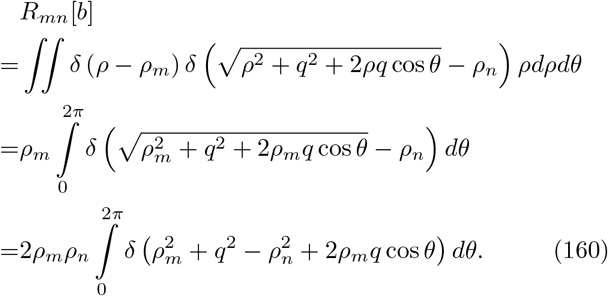

Given

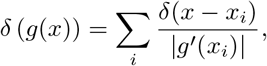

for all roots of *g*(*x*_*i*_) = 0; for 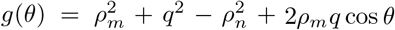, we have roots

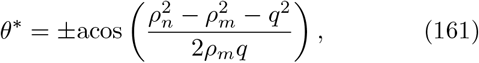

when *q* ∈ [*ρ*_*n*_ − *ρ*_*m*_, *ρ*_*n*_ + *ρ*_*m*_], otherwise *R*_*mn*_[*b*] = 0 when *g*(*θ*) = 0 has no roots. So, for both two *θ*^∗^, there are

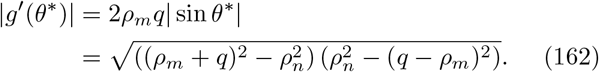

In the end we have:

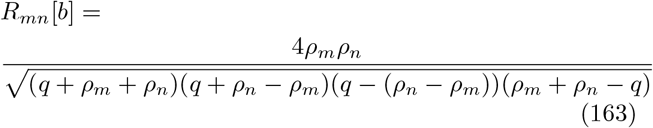

for *q* ∈ [*ρ*_*n*_ − *ρ*_*m*_, *ρ*_*n*_ + *ρ*_*m*_], otherwise *R*_*mn*_[*b*] = 0. For *R*_11_ and *R*_22_, there are two singularity points: *q* = 0 and *q* = 2*ρ*_1_ or *q* = 2*ρ*_2_. For *R*_12_[*b*], the singularity points are *q* = *ρ*_2_ − *ρ*_1_ and *q* = *ρ*_2_ + *ρ*_1_. For our parsimonious analysis, we need only consider *R*[*b*] = *R*_11_[*b*] + *R*_22_[*b*] + 2*R*_12_[*b*] nearby these five singularity points:

(a) Near *q* = 0,

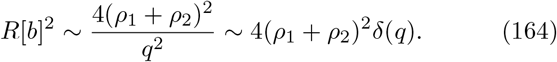

(b) Near *q* = 2*ρ*_1_, i.e. 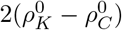:

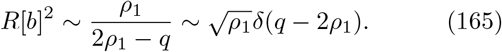

(c) Near *q* = 2*ρ*_2_, i.e. 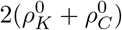:

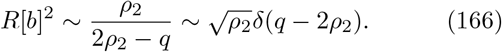

(d) Near *q* = *ρ*_2_ − *ρ*_1_, i.e. 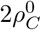:

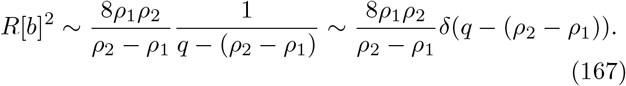

(e) Near *q* = *ρ*_2_ + *ρ*_1_, i.e. 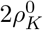:

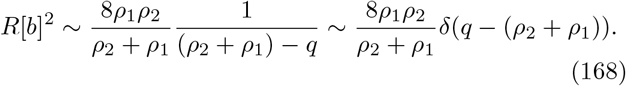

Finally, consider the inverse Hankel transformation:

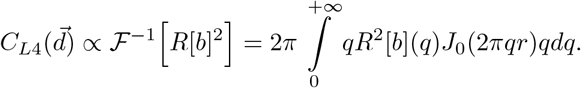

Substituting Eqs. (165) - (168) in the above (ignore Eq. (164) nearby *q* = 0 for low-frequency contents) yields:

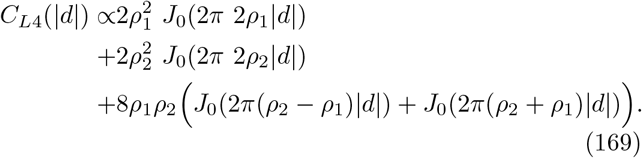

Here ignoring the relatively small term 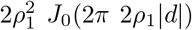 since 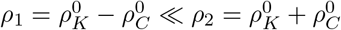, we get Eq. (47).

